# Peripheral linker mediates ACP’s recognition of DH and stabilizes Mycobacterium tuberculosis FAS-I

**DOI:** 10.1101/2023.06.27.546713

**Authors:** Akhil Kumar, Manisha Sharma, Harshwardhan H. Katkar

## Abstract

Incomplete structural details of Mycobacterium tuberculosis (Mtb) fatty acid synthase-I (FAS-I) at near-atomic resolution have limited our understanding of the shuttling mechanism of its mobile acyl carrier protein (ACP). Here, we have performed atomistic molecular dynamics simulation of Mtb FAS- I with a homology-modelled structure of ACP stalled at dehydratase (DH), and identified key residues that mediate anchoring of the recognition helix of ACP near DH. The observed distance between catalytic residues of ACP and DH agrees with that reported for fungal FAS-I. Further, the conformation of the peripheral linker is found to be crucial in stabilizing ACP near DH. Correlated inter-domain motion is observed between DH, enoyl reductase (ER) and malonyl/palmitoyl transferase (MPT); consistent with prior experimental reports of fungal and Mtb FAS-I.

## 2 Introduction

Mtb FAS-I is a multi-domain, multi-functional essential enzyme with a distinct capacity of *de novo* synthesis of long fatty acids^1–5^ and is a potential therapeutic target.^6, 7^ FAS-I is also found in fungi, mammals, and higher eukaryotes.^8–11^ Fungal and mycobacterial FAS-I have a structure resembling a closed barrel-shaped hexamer complex.^12, 13^ Despite these structural similarities, significant evolutionary gaps and mechanistic differences exist between them.^13, 14^ About 50% of amino acids are involved in scaffolding of fungal FAS-I, while only about 35% are used Mtb FAS-I. ^13^ Thus, Mtb FAS-I is a minimalist version of fungal FAS-I structure, lacking the entire phosphopantetheinyl transferase domain.^13, 14^ Fungal FAS-I has domains distributed across six identical α and six identical *β* chains.^15, 16^ However, in Mtb FAS-I, a pair of these α and *β* chains join to form a single chain containing all seven domains.^13^ Simply described, Mtb FAS-I structure has two domes (upper and lower) along with a central wheel. This closed arrangement is believed to facilitate coherent action of all enzymatic domains^17, 18^ and enable relatively higher local concentration of active sites, speculated to provide better control and selectivity.^19, 20^ Each dome consists of three identical chains with three distinct reaction chambers. Each chain consists of seven enzymatic domains along its primary sequence: acetyl transferase (AT), enoyl reductase (ER), dehydratase (DH), malonyl/ palmitoyl transferase (MPT), acyl carrier protein (ACP), ketoacyl reductase (KR), and ketoacyl synthase (KS).^13, 14^ Of these, ACP is a mobile domain that shuttles the growing fatty acid substrate between the remainder six catalytic domains to produce two classes of long-chain fatty acids (C_16_-C_18_ or C_24_-C_26_).^3, 5, 10, 21–25^ Gipson et al. reported ACP occupancy of approximately 30% near KS, ER, and AT, around 10% near KR, and less than 10% near MPT and DH in yeast FAS-I,^26^ suggesting that experimental determination of DH-ACP structure is challenging. However, the partial structure of fungal FAS-I with ACP halted at DH was recently resolved using cryogenic electron microscopy (cryo-EM).^17^

Research conducted in the past two decades has significantly enhanced our understanding of the structure and mechanism of fungal and bacterial FAS-I.^12, 14, 26–32^ However, detailed structural information about mycobacterial FAS-I was unavailable prior to two recent studies published in 2013 by Boehringer et al. and by Ciccarelli et al., which reported cryo-EM structures of *Mycobacterium smegmatis* and Mtb at 7.5 Å and 20 Å, respectively.^13, 14^ In all the reported cryo-EM structures of mycobacterial FAS-I till date, the structure of ACP was excluded, owing to the conformational flexibility of ACP, resulting in weak electron density. ACP shuttling is facilitated by two flexible linkers: the central linker (CL), connecting ACP with KR through the central wheel, and the peripheral linker (PL), connecting MPT with ACP.^18, 26^ These linkers were also not observed in any fungal or Mtb FAS-I structures due to their inherent flexibility. Gipson et al. reported two unre - solved electron densities in yeast FAS-I, each measuring 64 Å in length, and suggested that these could correspond to two distinct conformations of PL near ER.^26^ They also reported a propensity for helix formation in PL based on computational secondary structure prediction. Maier et al. suggested that higher proline and alanine content in PL could lead to increased rigidity, potentially limiting the entanglement and reducing the likelihood of free diffusion of each ACP to the nearest active sites in the reaction chamber of fungal FAS-I.^18^ A MARTINI coarse-grained simulation study suggested that in the absence of either PL or CL, the probability of ACP accessing the active site decreased by 70% and 30%, respectively, highlighting the importance of PL.^33^

It is known that ACP, and not the 4’-phosphopantetheine (PPT) arm that binds to it through a phosphodiester bond at a conserved serine residue^22^ in bacterial^23, 34^, animal^24^, and yeast^25^ FAS-I is responsible for shuttling of the substrate during the elongation.^10^ Fungal FAS-I ACP has a different structure than its bacterial counterpart *E. coli*, with four additional C-terminal alpha-helices forming a structural core.^31^ The four N-terminal helices in fungal ACP structurally align with bacterial and plant homologues, and constitute the catalytic core of ACP.^35–37^ The structural core has been reported to be more flexible than the catalytic core.^17^ Three of these catalytic core helices, α-1, α-2, and α-4, run roughly parallel to each other, while the remainder helix α-3 is shorter and nearly perpendicular to the other three. The precise role of the structural core of ACP is not fully understood. It is known to provide stability to the catalytic core and facilitate interaction via contributing to the surface area of the binding interface, while stalled at KS^28, 31, 38^ and AT.^39^ Lou et al. were able to resolve the surface binding between the catalytic core of ACP and ER at a relatively higher resolution compared to that between the structural core of ACP and ER using cryo-EM on fungal FAS-I. ^38^ To unravel the molecular intricacies of the interactions between the structural core and DH, it is imperative to conduct a thorough investigation into the DH-ACP interaction.

All-alpha helix configuration of ACP is believed to provide it with the flexibility necessary to bind acyl chains of varying lengths during the elongation phase.^31^ Furthermore, the catalytic core of ACP contains a dynamic pocket that can accommodate longer substrates, thereby protecting the expanding substrate linked to the PPT arm while shuttling between the domains.^35^ The residue SER180 between helix α-7 and α-8 in fungal FAS-I serves as the attachment site for the PPT arm.^40–43^ A highly conserved helix (α-2) succeeding SER180 aids ACP in recognizing other domains.^44–47^ In fungal and mammalian FAS-I, the distance between the PPT-binding site in ACP and the catalytic site in KS, KR, ER, and AT has been reported to be about 19 Å, 20 Å, 21 Å, and 18 Å, respectively.^26, 39^ This distance corresponds to the length of the PPT arm, where the substrate binds.^26, 31, 48^ The reactive site of the arm can extend and reach the active sites located in other domains. How-ever, the precise mechanism that enables ACP of Mtb FAS-I to reach the optimal distance between the substrate-binding site and the active sites of DH is still unknown.

FAS-I from *Saccharomyces cerevisiae* (SC) is often used as a representative example of fungal FAS-I. Its structure has been recently determined at near-atomic resolution, providing valuable information on this enzyme.^49^ The lack of structural information for the mycobacterial FAS-I complex at near-atomic resolution limited the determination of key interactions and dynamics at the domain and sub-domain scales, which can provide crucial insights for identifying novel Mtb drug targets. Thus, there is a need for a detailed investigation of Mtb FAS-I structure and dynamics, as was previously highlighted.^13, 14^ Recently, a partial nearly-atomic resolution structure of Mtb FAS-I was determined using cryo-EM.^50^ The structure was reported at a resolution of 3.3 Å, which included all the domains except for ACP, PL and CL, due to their weak electron density. However, Elad et al. have also reported observing an electron cloud with a relatively weak density in the vicinity of the KS, predicted to correspond to ACP.^50^

Our study utilizes computational tools to investigate the substrate shuttling mechanism of Mtb FAS-I at the dehydration step, where ACP stalls at DH. To accomplish this, we leverage two recent studies, cryo-EM structure of Mtb FAS-I at near-atomic resolution excluding ACP and linkers ^50^, and partial cryo-EM structure of fungal FAS-I with ACP stalled at DH.^17^ Our primary objective is to model the missing ACP and linkers in the former structure and use the latter to generate a stable conformation of the complex with ACP near DH, which is difficult to resolve using cryo-EM experiments due to its low occupancy at DH.^26^ We examine key residues involved in DH and ACP’s structural and catalytic core interactions. We investigated the role of specific residues in mediating the conserved residue SER1787 (SER180 in fungal, SER36 in *E. coli*) on ACP to reach an optimal distance from the catalytic center of DH for efficient catalysis, as well as the impact of the conformation of the linker on the stability of DH-ACP structure. Additionally, we investigate the coordinated motion of DH, MPT, and ER and observe ACP’s steric blockade of the neighbouring ER. This characterization provides insights into the dynamic interactions within the Mtb FAS-I complex.

## 3 Results

A homology model with ACP stalled at KS was generated using the partial resolved cryo-EM structure of Mtb FAS-I (PDB ID 6GJC)^50^, with fungal FAS-I ACP (PDB ID 6TA1) serving as a template^49^ (**see section 7.1, Figure 1A and 1D**). A reference structure of ACP stalled at DH was obtained by aligning DH of PDB ID 6WC7^17^ (fungal DH-ACP) with the above homology model and superimposing homology modelled Mtb FAS-I ACP on the resulting structure of fungal FAS-I ACP (**see section 7.3, Figure 1G**). Targeted molecular dynamics (TMD1) simulation was performed using this reference structure (**see section 7.3, Figure 1H**) and subsequentially with PL included in the reference structure TMD2 (**see section 7.4, Figure 1J and 1K**). Details are discussed in the STAR method section, and the results from the 50 ns (**Figure 1I**) and 100 ns (**Figure 1L**) classical molecular dynamics simulation (MD), referred as traj1_50ns_ (TMD1 without PL) and traj2_100ns_ (TMD2 with PL), respectively, are discussed in **section 3**.

**Figure 1.**
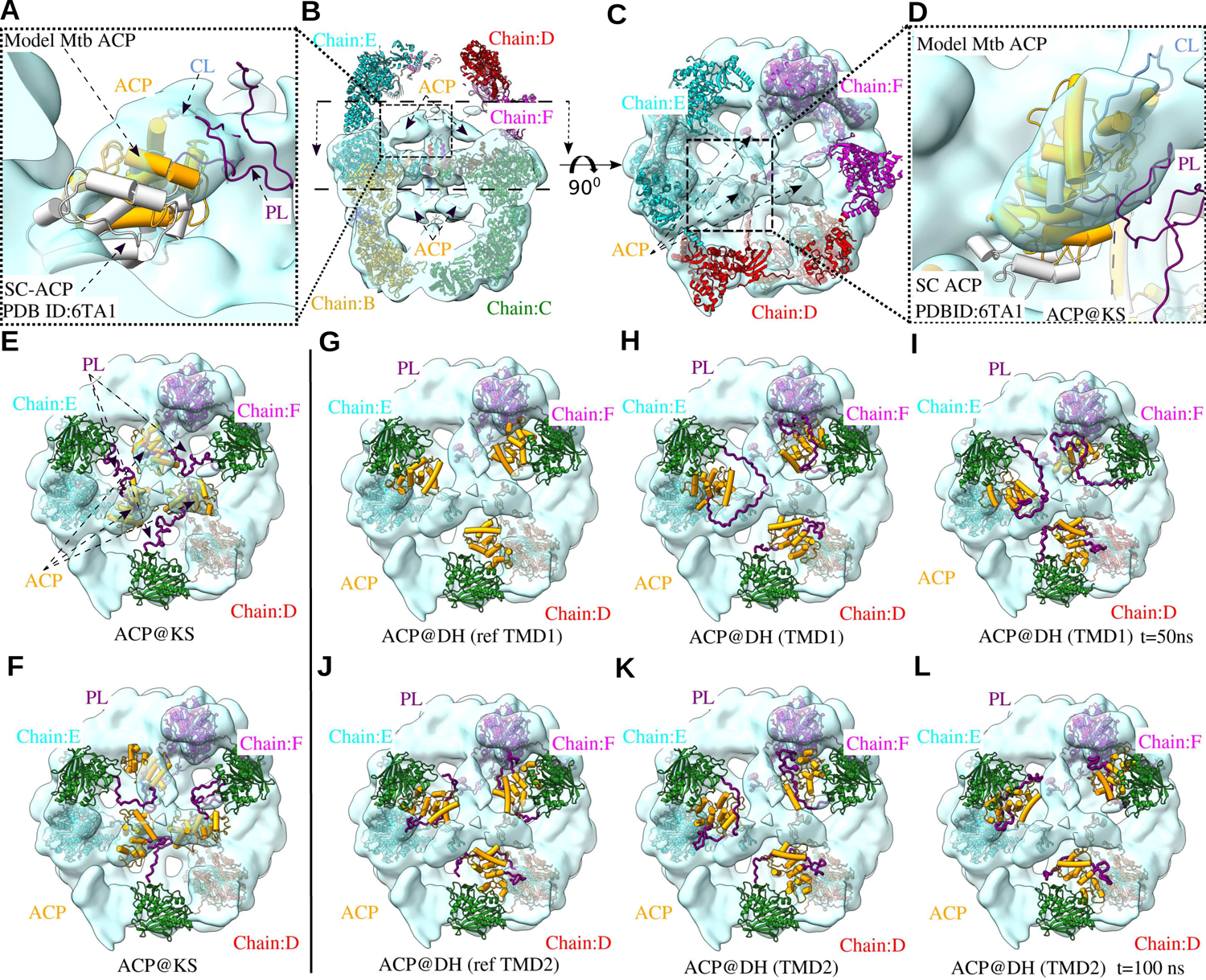
Positional characterization of ACP and PL in Mtb FAS-I complex through TMD simulations (see also Video S1). (A-D) The structures of SC and Mtb FAS-I, including the homology model of Mtb ACP; (A) and (D) show a focussed side and top view of weak electron density (putative location of ACP, shown as cyan clouds) reported near KS of chain E in Mtb FAS-I (PDB ID 6GJC), the superimposed structure of chain A from SC and the homology modelled and translated ACP of Mtb FAS-I (orange tubes). The predicted structures of CL and PL are shown as a purple cartoon. (B) and (C) show the side and top view of all six chains of Mtb FAS-I (cartoon) along with the corresponding electron density (EMD-0011).^50^ (E and F) The locations of homology-modelled structures of ACP stalled at KS and PL, relative to DH (green cartoon) in top dome of 6GJC and new conformation after equilibration. (G) The reference structure with modelled ACP at DH for TMD1. (H and I) Top view of Mtb FAS-I complex pdb (chains D, F, and F) depicting ACP stalled at DH and PL positions at the end of 10 ns TMD1 and traj1_50ns_, respectively. (J) Top view showing the reference pdb (chains D, F, and F) with ACP stalled at DH based on TMD1 chain D and PL have been included in TMD2 (absent in TMD1 (G). (K and L) Top view of Mtb FAS-I complex pdb (chains D, F, and F) with ACP stalled at DH, and PL indicating the position at the end of 10 ns TMD2 and traj2_100ns_, respectively.

### 3.1 Stable DH-ACP interaction in Chain D: Insights from Mtb FAS-I complex in traj1_50ns_

We monitored root mean square deviation (RMSD) and radius of gyration (R_g_) of the backbone atoms as indicators of the compactness and unfolding of each chain to assess the stability of the resulting homology modelled Mtb FAS-I complex with ACP stalled at DH. Each chain in traj1_50ns_ was aligned to its first frame using backbone atoms to remove rigid body translations and rotations before computing its RMSD or R_g_. **Figure 2A** shows the backbone RMSD of each of the six chains in the complex, revealing an increase in the RMSD values for all chains over time. In the last 25 ns of the traj1_50ns_, the time-averaged backbone RMSD of chains was between 5.4 Å and 6.9 Å. **Figure 2C** shows that the backbone R_g_ of each of the six chains does not show a prominent increase, except for chain A. Thus, each chain undergoes internal rearrangements, dominated by large fluctuations in CL and PL regions (data not shown), which increase their RMSD with time while their overall shape does not change significantly. To test whether ACP persistently stalls near DH in traj1_50ns_, the center of mass (COM) distance of DH and ACP (COM_DH-ACP_) was computed for all six chains (**Figure 2E**). In chain D, this distance gradually decreased during the simulation, suggesting that, unlike in all other chains, a persistent interaction was likely established between DH and ACP of chain D in the traj1_50ns_. The RMSD of PL in each chain, calculated after aligning the backbone atoms in the neighbouring MPT, DH, and ACP, was found to be stable in chain D (**Figure 7A**). Chains D and E exhibited a unique conformation of PL away from all the domains, wrapped across ACP. In the rest of the chains, PL acquires a conformation near KS, ER and DH (**Video S2**). Thus, for refining the homology-modelled Mtb FAS-I complex with ACP stalled at DH, the conformation of chain D at the end of the traj1_50ns_ was used to generate a new reference structure inclusive of ACP and PL to perform another targeted MD simulation (TMD2). TMD2 was followed by slow equilibration and 100 ns production run (traj2_100ns_) (**see section 7.4**).

**Figure 2.**
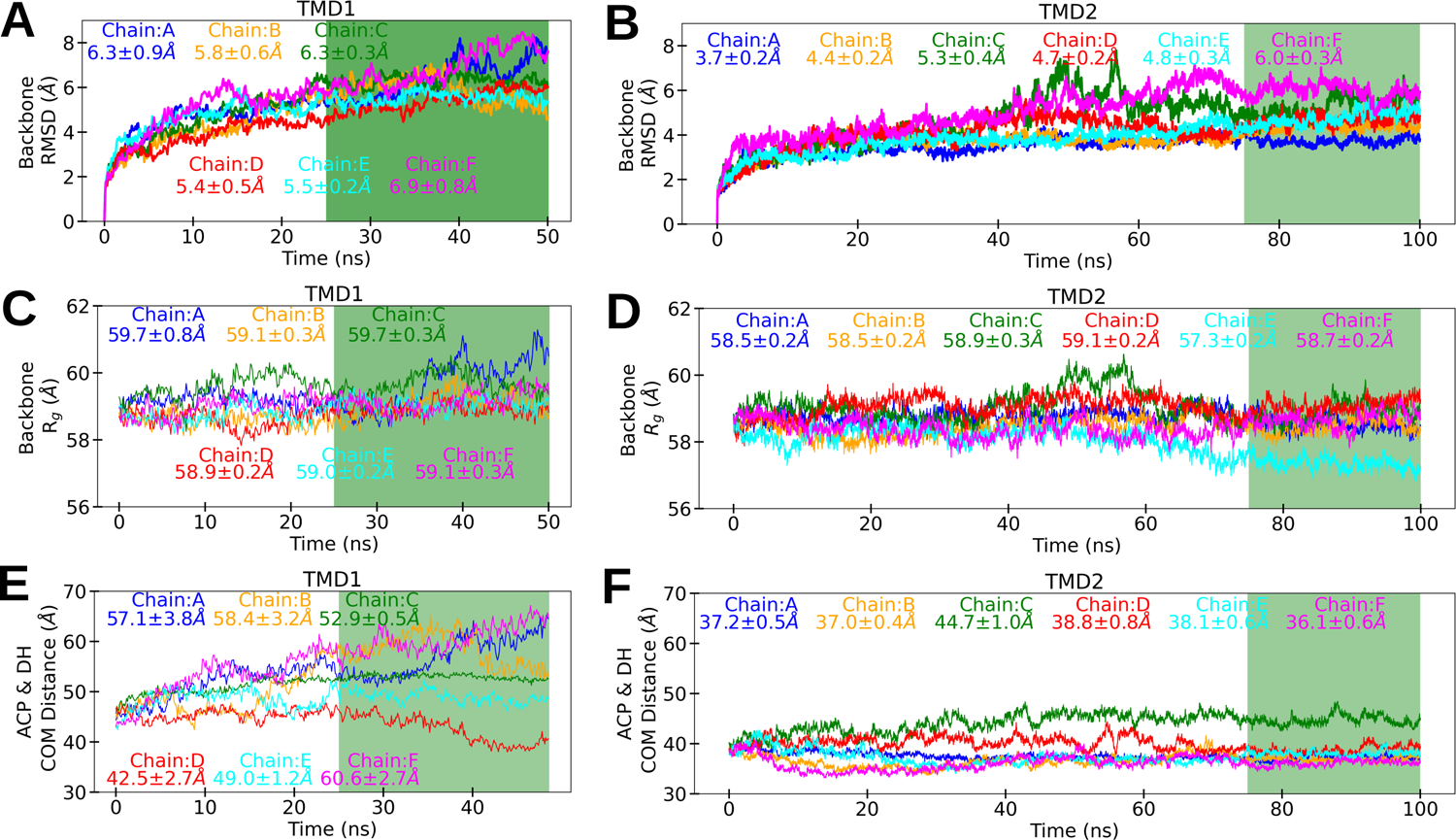
Comparison between traj1_50ns_ with traj2_100ns_. (A and B) RMSD of backbone atoms in each chain as a function of time, indicating lower RMSD in traj2_100ns_ compared to traj1_50ns_ for all six chains (The legend shows mean and standard deviation in the last 25 ns (green region) in A-F). (C and D) R_g_ of backbone atoms in each chain as a function of time, indicating R_g_ is relatively lower for all six chains in traj2_100ns_. (E and F) COM_DH-ACP_ distance as a function of time. In contrast to other chains, COM_DH-ACP_ distance of chain D decreases in the traj1_50ns_, while it remains consistently low in all six chains during traj2_100ns_.

### 3.2 Improved stability of Mtb FAS-I complex in traj2_100ns_

The conformation of Mtb FAS-I complex obtained after TMD2 showed significant improvement in structural stability compared to that obtained after TMD1. A side-by-side comparison of RMSD and R_g_ of backbone atoms and COM_DH-ACP_ distance in traj1_50ns_ and traj2_100ns_ are shown in **Figure 2**. After about 10 ns of traj2_100ns_, RMSD of individual chains were found to be stable. In the last 25 ns of traj2_100ns_, the time-averaged backbone RMSD of chains was between 3.7 Å and 6.0 Å (**Figure 2B**). The R_g_ value was relatively stable for all six chains traj2_100ns_, with R_g_ of chain E decreasing during the simulation (**Figure 2D**). Remarkably with the exception of chain C, the COM_DH-ACP_ distance in the traj2_100ns_ remained low relative to traj1_50ns_. Although an increase in COM_DH-ACP_ was observed in chain C, it was less prominent compared to that in traj1_50ns_ (**Figure 2E and 2F**). The relatively low backbone RMSD and R_g_, along with nearly-constant COM_DH-ACP_ distance in traj2_100ns_ indicate persistent contact between DH-ACP in all six chains of the complex. The conformation of PL in all six chains was similar to the unique conformation of chain D in traj1_50ns_.

After ensuring the overall stability of the complex, the residue-level dynamics during the traj2_100ns_ were investigated using the root-mean-square fluctuations (RMSF) of Cα atoms in each chain with respect to the initial conformation (**Figure 3C**). Residues belonging to CL are the most flexible in traj2_100ns_ (**Figure 3E**), with a six-chain-average RMSF of 6.57 Å (**Table S1**). ACP and PL are also observed to be flexible (**Figure 3B and 3D**), with average RMSF values of 4.71 Å and 3.38 Å, respectively. Both PL and CL lack a rigid secondary structure, which is mechanistically necessary to enable the shuttling of ACP in the reaction chamber.^12, 33^ Within ACP, the first four helices (catalytic core) are observed to have a lower RMSF relative to the other four helices (structural core) (**Figure 3B**). **Figure 3A** shows fluctuations in DH catalytic and surface residues. As characterized in **section 3.4.a**, residue-level interactions are established between the catalytic core and DH. Thus, the catalytic core is expected to have a lower RMSF when ACP is stalled at DH. The lower RMSF in the catalytic core of Mtb ACP observed in our simulation has also been observed in the experimentally determined structures of fungal FAS-I.^17, 38^ The catalytic core of fungal ACP was reported at a resolution of 5 Å, while the structural core was resolved at ∼9 Å, suggesting that the catalytic core is less flexible than the structural core^17^.

**Figure 3.**
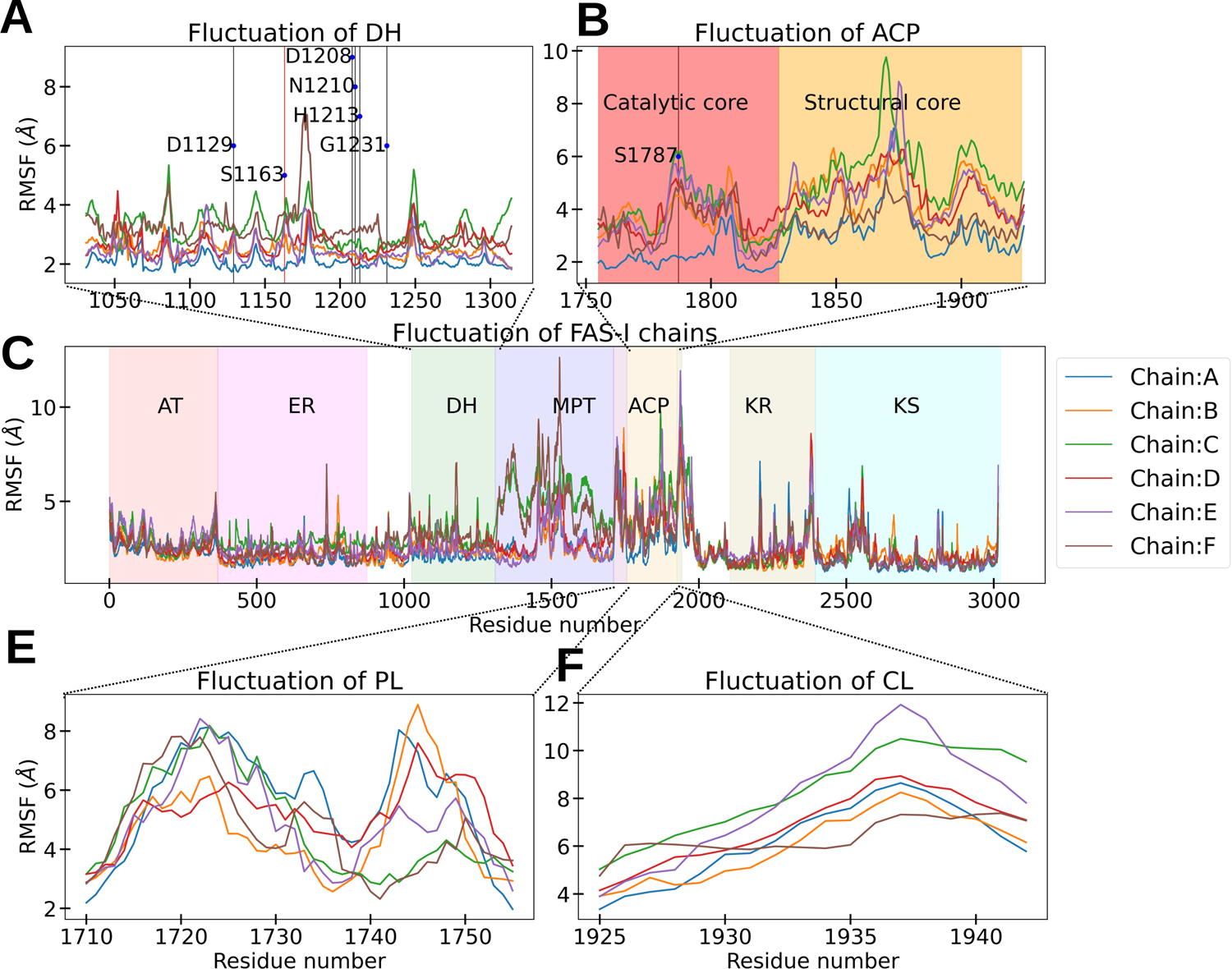
Residue-level fluctuation of Mtb FAS-I complex in traj2_100ns_. (A**-**E) RMSF of Cα atoms after aligning to its first frame for all six chains of Mtb FAS-I complex, different color span shows seven domains in each chain and subplots shows RMSF of (A) DH (B) ACP’s catalytic and structural core (C) whole complex (D) PL and (E) CL.

In our analysis of RMSF, MPT showed a high degree of flexibility, with an average RMSF of 2.61 Å. Notably, residues between VAL1451 to PHE1498 in MPT exhibited higher fluctuation, with an RMSF of 3.72 Å (**Table S1, Figure 3C**). Gipson et al. have reported conformational differences in the reaction chamber between x-ray and cryo-EM structure and suggested that in cryo-EM experimental structure, an inward movement of MPT and DH with simultaneous outward movement of ER could result from crystal contacts that involve the loop of MPT.^26^ Ciccarelli et al. also demonstrated that MPT in Mtb FAS-I exhibits conformational variability. They observed three different conformations of MPT, facilitated by the associated movement of DH such that together they appear to pivot around the ER.^13^ Similar movement is observed in traj2_100ns_. A positive correlation between the COM distance of KR-MPT and KR-DH is observed, suggesting that these two domains move in phase **(Figure S1A)**. A negative correlation was observed in the COM distance between ER-KR and KR-MPT **(Figure S1B)**, suggesting that when ER moves towards KR, MPT simultaneously moves away from KR.

### 3.3 Internal stability and catalytic core dynamics of ACP

The secondary structure of our homology-modelled ACP consisted of 168 residues (from ALA1756 to GLY1924) participating in the formation of eight α-helices. We monitored the backbone RMSD and R_g_ of ACP of each chain in the traj2_100ns_ after aligning the backbone atoms of each ACP in the trajectory to its first frame. The average RMSD during the last 25 ns of the simulation was steady and varied between 3.6 Å and 4.2 Å across all six chains (**Figure 4A**). The average R_g_ of all six chains over this duration varied between 16.1 Å and 17.0 Å across all six chains (**Figure 4B**). Thus, the backbone structure of ACP did not undergo significant changes in the traj2_100ns_. The average fraction of native contacts steadily remained above 88 % in the last 25 ns of the simulation, indicating stable intra-domain interactions resulting in a stable structure of Mtb ACP (**Figure 4C**)

**Figure 4.**
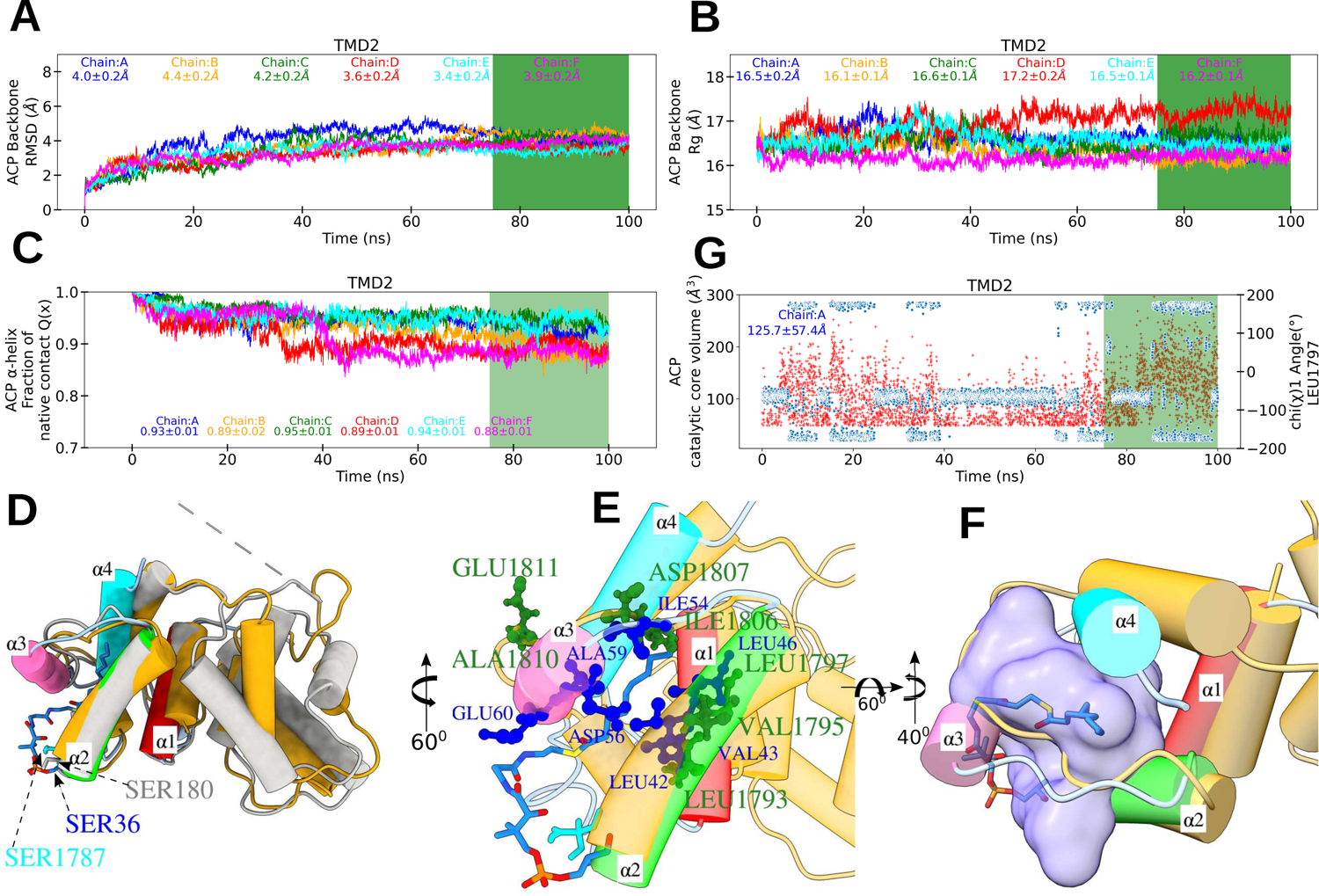
ACP: structural integrity, recognition helix, and binding pocket dynamics. (A, B, and C) RMSD, R_g_ for backbone atom and the fraction of native contacts for eight α-helix were calculated after trajectories aligned using backbone atoms of respective chains of ACP in traj2_100ns_, mean and standard deviation were calculated for the last 25 ns (green region) reported in each sub-figure legend, indicating ACP structural integrity (**see also Video S3)**. (D) Average structure of Mtb ACP in last 25 ns for all chains (orange cartoon) superimposed with the crystal structure of *E. coli* decanoyl-FAS-II ACP (PDB ID2FAE) and SC ACP (PDB ID 6TA1) used as a template for homology modelling. Three of four α-helix in Mtb ACP resemble α-1, α-2 and α-4 helices from *E. coli* FAS-II ACP. α-3 helix is reported to switch between loop helix conformation^56, 57^ and showed loop structure in 6TA1. α-2 and α-8 are recognition helices in *E. coli* and SC, respectively. The corresponding helix has been identified and labelled as α-2. SER, the PPT attachment residue, is present in all species highlighted in different color stick representations indicated via black arrow. (E) The catalytic core of *E. coli* FAS-II ACP bound with decanoyl, as reported by Roujeinikova et al.^35^, highlights key aliphatic residues inside the hydrophobic pocket (blue stick representation), as demonstrated by Chan et al..^58^ Notably, a similar arrangement of aliphatic residues on the α-2 helix of Mtb ACP model was observed to sequester the acyl chain (green ball and stick representation). (F) The catalytic core pocket was detected using trj_cavity^62^ and represented as a purple surface in a transparent representation of the average structure of Mtb ACP for the last 25 ns for all chains. In addition, the bound decanoyl from *E. coli* from PDB ID 2FAE was sequestered in the predicted pocket. (G) Catalytic core cavity volume as a function of time indicates that the pocket in ACP catalytic core is highly dynamic.

The residues from SER1787 to THR1827 constitute the catalytic core region of ACP, which is highly conserved across several species.^35, 37, 51–53^ In addition to four α-helices of the catalytic core region which are also present in *E. coli* FAS-II, Mtb ACP has four additional helices.^31, 35^ By aligning the average structure of the last 25 ns for all six chains with crystal structures of *E. coli* FAS-II ACP, and SC ACP, we have identified the recognition helix in Mtb ACP (**Video S3, Figure 4D**), which is labelled as α-2 helix in *E. coli* FAS-II ACP and α-8 helix in SC ACP.^31, 35^ This helix is reported to be responsible for recognizing of other domains in the FAS-I complex. ^35, 46, 47, 54, 55^ The helices labelled α-1, α-2, and α-4 in *E. coli* ACP directly map to the three remaining helices of the catalytic core of Mtb ACP (α-1, α-3, α-4). The α-3 helix, reported as a short helix in *E. coli* FAS-II ACP, adopts a loop conformation in our homology model of Mtb ACP, resembling SC ACP. The α*-* 3 helix is known to be flexible and does not have a conserved fold in *Plasmodium falciparum* FAS-II ACPs.^56, 57^ This helix was reported to dynamically switch between loop and helix conformation in the molecular dynamics simulation of *Plasmodium falciparum* ACP.^56^ We have computed the secondary structure of the catalytic core for the last 25 ns of traj2_100ns_ for all chains (**Figure S2A**). Our simulation trajectory shows similar results in which α*-*3 transition between loop and helix conformation **(Video S3)**.

The hydrophobic binding pocket in *E. coli* FAS-II ACP is formed by the combination of α-2, α-3, α-4 helices, and α-1, α-2, α-4 helices, consisting of 16 different residues.^58^ In Mtb ACP, we have identified three (LEU1793, VAL1794, and LEU1797) of four residues along the recognition helix, which are known to form the hydrophobic binding pocket in *B. subtilis* (VAL39, LEU42, VAL43, and LEU46) and *E. coli* FAS-II ACPs (VAL40, LEU42, VAL43, and LEU46).^53, 58^ Additionally, residues ASP1807, ALA1810, and GLU1811 from the α-3 helix of Mtb ACP have been identified as part of the pocket, corresponding to ASP56, ALA59, and GLU60 in the short α-3 helix of *E. coli* FAS-II ACP. Furthermore, ILE1762 in Mtb FAS-I corresponds to the common residue ILE11 in *E. coli* FAS-II, which connects α-2 to α-3. Thus, we have identified seven common residues out of sixteen hydrophobic binding pocket-forming residues in *E. coli* FAS-II ACP and Mtb ACP (**Figure 4E**).

Throughout the simulation, the binding pocket (**Figure 4F**) exhibited significant variability in size, ranging from 49.39 Å^3^ to 296.35 Å^3.^. The dynamics of the binding pocket in chain A of Mtb ACP were investigated by measuring its volume. We categorize it as either open or closed, assuming a threshold value of 50 Å^3^. Approximately 29% of the simulation time, the pocket was closed. The average size of the pocket fluctuated during the simulation, with a volume of 113.34 Å^3^ in the first 20 ns, 95.52 Å^3^ between 20 ns and 40 ns, 78.84 Å^3^ between 40 ns to 60 ns, 91.52 Å^3^ between 60 ns to 80 ns, 141.81 Å^3^ from 80 ns to 100 ns (**Figure 4G red dots**). A similar trend was observed in a MD simulation study of *E. coli* FAS-II ACP, where the binding pocket expanded slightly with smaller acyl groups and reached a maximum size of 231.9 Å^3^ in the presence of decanoyl substrate.^58^ The apo pocket size was also large, reaching approximately 115 Å^3^. These findings suggest that Mtb apo binding pockets are also dynamic and can expand to accommodate longer substrates. MD simulation of *E. coli* FAS-II ACP, revealed that the movement of side chains of LEU42 and LEU46 controlled the opening and closing of the two sub-pockets.^58^ We measured the angle made by the sidechain with the backbone for corresponding residues LEU1793 and LEU1797 in Mtb FAS-I during traj2_100ns_. The binding pocket volume increase is always accompanied by LEU1797 side-chain making an angle of less than −100° or greater than 100° with its backbone (**Figure 4G blue dots**).

To investigate the interactions among the catalytic core residues that may contribute to its stability, we conducted network analysis using RING 3.0^59^ for the last 25 ns of the traj2_100ns_ for all six chains using Cα atoms (**Figure S2B**). Hydrophobic residues dominate the catalytic core of ACP, with ILE1762, ILE1779, and LEU1817 exhibiting the highest degree of interaction. The recognition helix residues LEU1793, LEU1794, and LEU1797 form a dense interaction network. The high degree of interaction among hydrophobic residues suggests that hydrophobic interactions are crucial for maintaining the secondary structure of the catalytic core of ACP. This observation is consistent with previous studies on FAS-II ACP, which have shown that hydrophobic interactions are responsible for maintaining stable folds despite differences in primary sequences across different species.^52, 55^ A site-directed mutagenesis study of ACP in Vibiro harveyi identified PHE50, ILE54, and ALA59 as essential residues for establishing interactions, crucial for the native conformation of ACP.^60^ ILE1806 and ALA1810, two of the corresponding residues in Mtb ACP, also show significant number of interactions in the network analysis. However, Mtb ACP lacks a corresponding PHE residue, which is essential for the stability of *E. coli* FAS-II ACP structure.^60^ Another important contact that stabilizes the helical bundle in E. coli and Plasmodium falciparum FAS-II ACPs is the VAL-ALA contact between VAL43 of helix α-2 and ALA68 of helix α-4 in E. coli or VAL41 of helix α-2 and ALA60 of helix α-3 in Plasmodium falciparum, respectively.^56, 58^ We have identified corresponding residues as VAL1795 and ALA1810 in Mtb ACP. The distance between these two residues fluctuates greatly (**Figure S2C**). As earlier MD simulation studies of *Plasmodium falciparum* ACP reported that, the distance between these two residues increases and acts as a gate opening for substrate delivery.^56^

### 3.4 Key Interactions of ACP with other domains

#### 3.4.a ACP stalled at DH and sterically blocks ER

After aligning DH of each chains, we examined DH-ACP interaction in traj2_100ns_ to highlight key interacting residues. We observed that ACP in chain A remained stable throughout the simulation (**Figure 5B**). In contrast, the recognition helix in chains B, D, and E first retracted from DH and then moved towards it to establish a stable conformation. Chains C and F did not show similar stable conformation. To determine which specific residues in DH and ACP are responsible for their stable anchoring, we examined the persistent contacts between DH-ACP for all six chains. The result revealed that residues SER 1787 and SER 1788 of ACP catalytic core interact with DH surface residues ASP1129, SER1163, ALA1164, GLU1165 (**Figure 5C**). In chain A, these interactions are present for more than 90% of the time (**Figure S3A**), whereas, in chain B and D, it forms after ∼60 ns (**Figure S3B and S3D**); for chain E and F, only ASP1129 and GLU1165 are formed but not other two residues (**Figure S3E and S3F**). For chain C, initially, GLU1165 and ASP1129 are in contact with SER1787 and SER 1788 for 20 ns. However, after 30 ns, the contact is broken and is not formed during the rest of the simulation (**Figure S3C)**. The increasing COM_DH-ACP_ distance in chain C also supported this trend (**Figure 2F**). In addtion to ACP’s catalytic core, we also observed the interaction of structural core residues GLU1899, GLY1900 and ALA1901 of chain A, which persistently interacted with DH residues ASP1182, THR1183, PRO1184, ARG1185 (**Figure 5A**).

**Figure 5.**
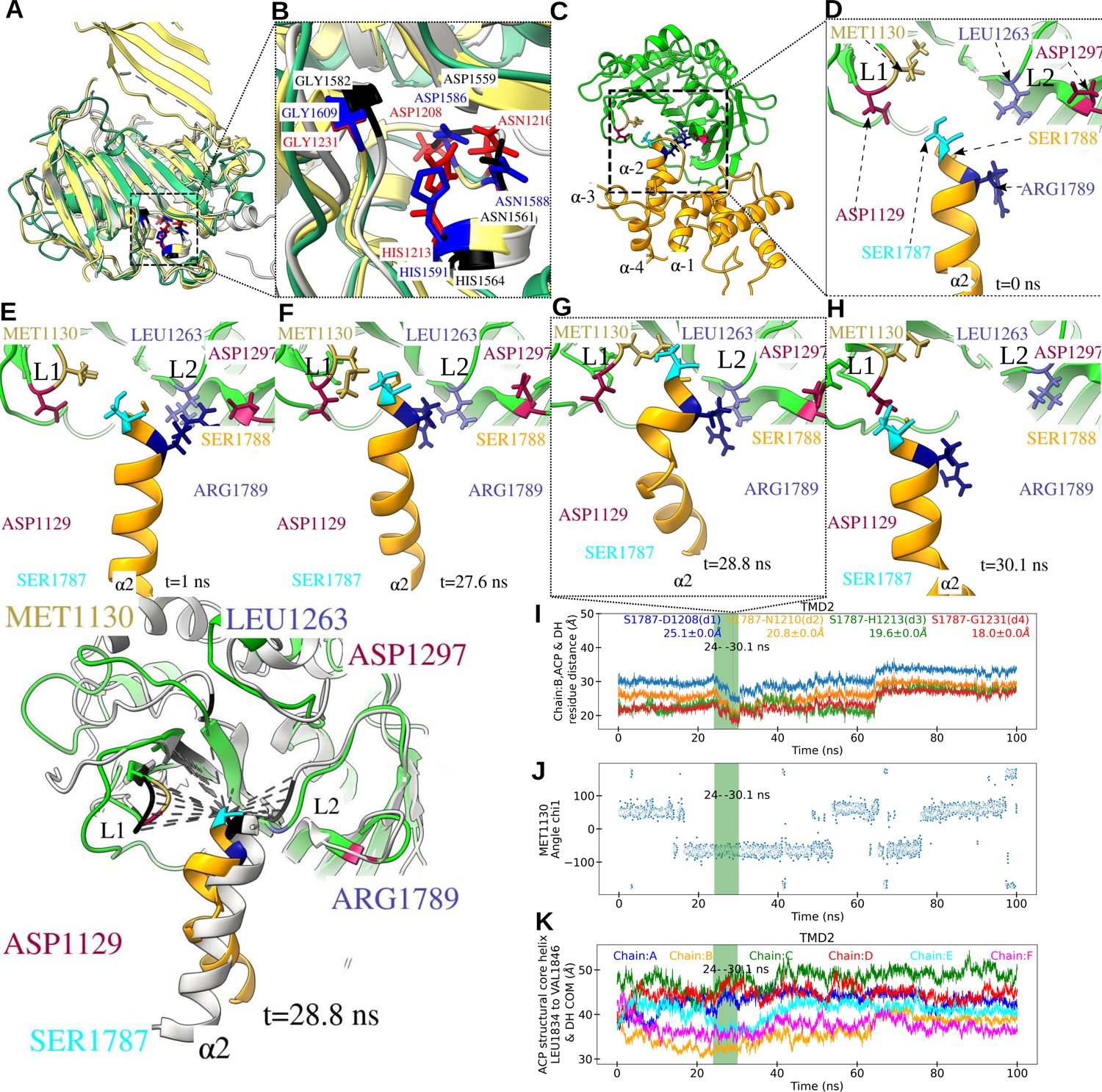
Insights into the interactions between the catalytic and structural core of ACP with DH. (A-C) Average structure of DH-ACP and key interacting residues. (B) Average structure of ACP (orange cartoon) and DH (green cartoon) for the last 25 ns. (A) Zoomed-in view of interacting amino acid residues between ACP structural core (GLU1899, GLY1900, ALA1901) and DH surface residues (THR1183, PRO1184, ARG1185), and (C) Zoomed-in view of the interaction between ACP recognition helix and surface residues of DH. SER1787 (cyan) and SER1788 (orange) of ACP are shown in stick representation, whereas DH residues (ASP1129, SER1163, ALA1164, GLU1165) are represented as stick in different color. (D) Throughout the simulation, ACP stalled at DH was observed to block the adjacent ER (magenta). (E) Zoomed-in view of ACP interacting with adjacent ER, highlighting key residues from α-2 and α-4 helices.

In our trajectory, while ACP is at DH, its catalytic core occupies volume around ER and sterically blocks it. We observed persistent interaction between ACP residue SER1787 with ER residue LYS478. Additionally, ACP residues SER1812, ASP1813, LEU1814, ALA1815, GLY1816 in persistent contact with ER residue VAL432 THR433, TYR464, LEU468, LEU818, GLY819. We also observed persistent contact between ACP LEU1776, ASP1777, SER1778 with ER’s TYR464 **(Figure 5D and 5E)**. A similar conclusion was drawn in a previous experimental study on fungi, where it was observed that while ACP stalled at DH, ER sterically blocked.^47^ The recognition helix, which plays a crucial role in recognizing the binding domain during the reaction cycle, is the primary source of these interactions. Interactions between ACP’s recognition helix and ER observed in tra-j2_100ns_ agree with the previous studies, have shown that after the dehydration step, ACP moves to ER for reduction, facilitated by the recognition helix, which requires minimal rearrangement from DH.^17^

#### 3.4.b Interaction of ACP’s recognition helix with DH

DH facilitates the dehydration step in fatty acid synthesis through acid/base pair residues that are conserved across different species.^61^ Catalytic residues involved in the reaction mechanism on DH have been identified in fungal FAS-I, including SC’s residues ASP1586, ASN1588, HIS1591, and GLY1609, as well as in *Thermomyces lanuginosus*, consisting of ASP1559, ASN 1561, HIS1564, and GLY1582.^27, 31^ By superimposing DH of SC and *Thermomyces lanuginosus* onto DH of Mtb, we were able to map the catalytic residues on Mtb DH as ASP1208, ASN1210, HIS1213 and GLY1231, (**Figure 6A and 6B**). We monitored four distances between, Cβ of ACP SER1787 and DH ASP1208 (CG), ASN1210(CA), HIS1213(NG1), and GLY1231(CA) for all six chains **(Figure S4 A-E and Figure 6I**), which we referre to as d1, d2, d3, and d4, respectively, in chain B (**Figure 6C, Video S4)**. **Figure 6I** shows at the beginning of the simulation, the distances (d1-d4) were 29.18 Å, 29.11 Å, 22.06 Å, and 20.42 Å, respectively. After 24 ns, all four distances exhibited a significant decrease. Distance d1, d2, d3 and d4 reach 25.05 Å, 20.77 Å, 19.55 Å, and 17.99 Å, respectively at 28.8 ns. The distances (d1-d4) were found to be at their minimum values (23.5 Å, 18.8 Å, 16.5 Å, 16.7 Å, respectively) at 30.1 ns after which, the distances gradually increased and reached close to their initial values at around 65 ns. Thereafter, they continued to increase and eventually remained stable throughout the simulation. A similar pattern was observed in the corresponding COM_DH-ACP_ distance (**Figure 2F).** This distance first decreased for the first 20 ns for chains B and F, then remained stable until 40 ns, after which it started to increase and stabilized after 70 ns. For the reaction to occur, the growing acyl chain attached to the PPT arm of ACP needs to reach the catalytic residues. Gipson et al. and Leibundgut et al. reported the optimal distance between ACP, SER180 (in yeast), where the PPT arm is expected to attach, and the catalytic domain of KS, KR, ER, and DH to be between 18 Å to 21 Å, corresponding to the length of the PPT arm.^26, 27^ We also found these distances in this range during our simulation, between 28.8 ns to 30.1 ns. (**Video S4**).

**Figure 6.**
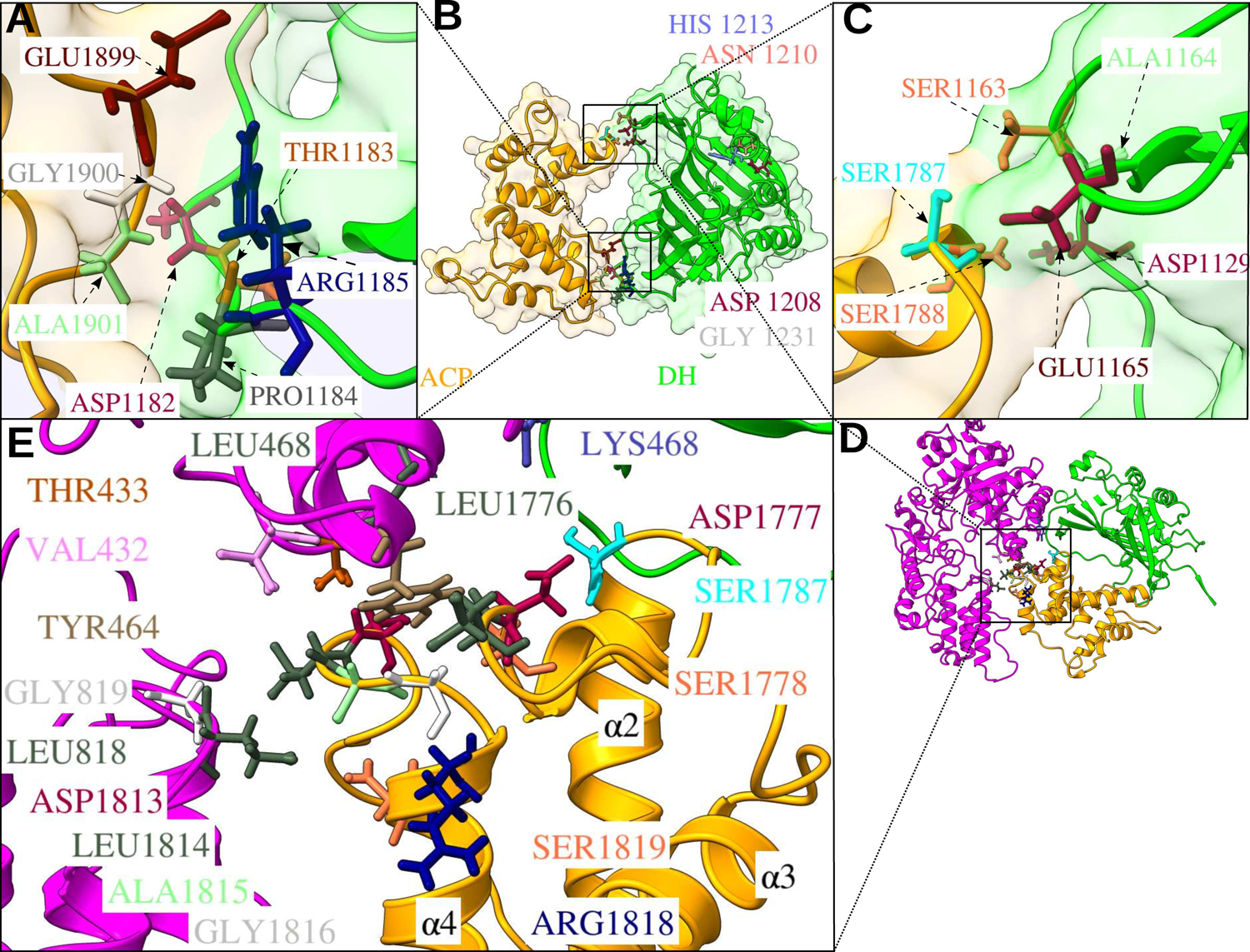
Exploring interactions between ACP recognition helix and DH. (A and B) Alignment of fungal DH (*Thermomyces lanuginosus i*n yellow cartoon and SC in gray cartoon) to Mtb DH (green cartoon) to identify corresponding catalytic residues in Mtb DH. (B) Zoomed-in view on catalytic residues (represented as stick) of *Thermomyces lanuginosus* (blue), SC (black), and Mtb DH (red). (C) Initial conformation of ACP (orange cartoon) stalled at DH (green cartoon), and all four catalytic core helices (α-1, α-2, α-3, α-4) are labelled. (D and H) Zoomed-in view of ACP recognition helix and DH, with the important interacting residues depicted in stick representation at different time intervals. Snapshots of the recognition helix at various time intervals showing the contribution of DH residues (LEU1297 and LEU1263, ASP1129, MET1130) and ACP residues (ARG1789, SER1788) to bring SER1787 into range for catalysis by DH. (I) Distances between DH catalytic residues (CG of ASP1208, NG1 of HIS1213, Cα of ASN1210, Cα of GLY1231) and ACP (Cβ of SER1787) d1,d2,d3 and d4 respectively are shown at 28.8ns. These distances observed at 28.8 ns agrees with previous results (shaded region shows event-1 window in I-J) (J) Change of MET1130 chi1 angle throughout the simulation time showing MET1130 assists SER1787 to reach the catalytic site to DH. (K) Analysis of COM distance shown for all chains, reveals that only chain B exhibited a shift towards DH from helix LEU1835 to VAL1846, suggesting an interaction between ACP structural core and DH. (L) Comparison of residues within 10 Å distance of SER180 (interactions are highlighted with dash lines) in SC DH-ACP structure (PDB ID 6WC7) with Mtb ACP’s recognition helix near DH at 28.8 ns.

After analyzing the trajectories, we found a series of interactions (**event-1**) between the residues of loops L1 (ASP1129, MET1130) and L2 (LEU1263) on DH with ACP’s recognition helix (**Figure 6D-6H**). Specifically, SER1787, SER1788, ARG1789 on ACP’s recognition helix and ASP1129, MET1130, LEU1263, ASP1297 on DH participate in the following series of events in order to bring ACP’s recognition helix closer to DH that led to the structure at 28.8 ns (**Figure 6G**). Firstly, ARG1789 forms a hydrogen bond with ASP1297 (**Figure 6E, t=1 ns)**. Subsequently, this bond breaks and ARG1789 forms a hydrogen bond with LEU1263 (**Video S4, t=24 ns)**. MET1130 pivots as shown in **Figure 6J** and forms a hydrogen bond with ASP1129 and SER1788 simultaneously (**Video S4, t=24.96 ns)**. Finally, MET1130 forms a hydrogen bond with SER1787 (**Figure 6F, Video S4, t=27.6 ns**) and drags it towards the catalytic residues of DH (**Figure 6G, t=28.8 ns**). After 30.1 ns, the recognition helix retracts and establishes interactions with ASP1129, which persisted throughout the simulation. The structure at 28.8 ns is superimposed with the structure of fungal DH-ACP. In the fungal FAS-I ACP, the interaction of SER180 (corresponding to SER1787 in Mtb) within 10 Å is shown in **Figure 6L**. We observed that the conformation of the recognition helix and surface residues of fungal DH, PRO1385, THR1386, VAL1613 (corresponding to L1 and L2 in Mtb), are similar to what we observed in traj2_100ns_ at 28.8 ns (**Figure 6L**).

#### 3.4.c Interaction of ACP’s structural core with DH

We investigated the influence of ACP’s structural core on DH-ACP interaction by computing the COM distance between them (results not shown). Our results indicated chains B and F had low COM distance between 10 ns to 35 ns. Further examination of the trajectories revealed that the shift of helix LEU1835 to VAL1846 towards DH is most prominent in chain B between 28-30 ns (**Figure 6K**). This shift was due to the movement of ARG1844’s side chain towards ARG1187 **(event-2)**. To compare this interaction in chain A and B, we examined the distance between these residues at different time instances during event-1. Initially, this distance in chain A and B is 16.55 Å and 10.33 Å, respectively. Interestingly, this distance gradually decreased (5.5 Å at t =24 ns, 4.8 Å at t=24.96 ns, 4.37 Å at t=27.5 ns) and reached a minimum (3.91 Å) at t=28.8 ns in chain B. In chain A the distance at these time instances was always greater than its initial value, i.e. 16.54 Å. After 30.1, this distance increased in both chains (**Video S5**). These observations suggest that the structural core plays a role in stabilizing the interaction between ACP and DH. However, these interactions subsequently broke and did not reform. This finding is consistent with previous studies that have shown the role of the structural core in promoting the interaction between two domains when ACP is situated near KS or AT.^28, 31, 38, 39^

#### 3.4.d Linker’s role in ACP-DH stability

RMSD of PL for each chain was calculated after aligning MPT, DH and ACP to the first frame of the traj2_100ns_. The average RMSD of PL for all six chains in traj2_100ns_ (7.15 Å) indicated higher stability than traj1_50ns_ (10.9 Å), as indicated in **Figure 7A and 7B**. In traj2_100ns_, after 75 ns, PL in all six chains showed RMSD in a range of 6.2 Å to 8.1 Å. As discussed in **section 3.1**, COM_DH-ACP_ distances exhibited stability in traj2_100ns_ in contrast to traj1_50ns_, plausibly due to completely different conformation of PL in the reaction chamber (similar to that in chain D of traj1_50ns_). Moreover, in traj2_100ns_, the linker of chain B adsorbed on to ACP. The distances measured from PL residue PRO1715 to ALA1726, ALA1726 to PRO1715, and ALA1726 to the COM of ACP (**Figure 7D**) indicated that the linker in chain B moved towards ACP while distance d1-d4 are in catalytic range (event-1) (**Video S5**). This observation highlights the significant role of the PL in facilitating the interaction between DH and ACP. However, we did not find any direct interaction between PL and DH. Thus, we have used a coarse-grained HENM model to validate PL interactions. Based on our analysis, the average spring constant between different domains was higher in traj2_100ns_ compared to traj1_50ns_, indicating stronger interactions in the traj2_100ns_. We also observed an improvement in PL-DH and DH-ACP interactions in traj2_100ns_ (**Figure 7C**). This positioning of the PL provides significant advantage as it efficiently inhibits any unfavourable interactions between the linker and the growing fatty acid substrate. However, we did not find any pronounced interaction with CL. Similar conclusions have been made earlier suggesting that PL contributions are more important than CL.^33^

**Figure 7.**
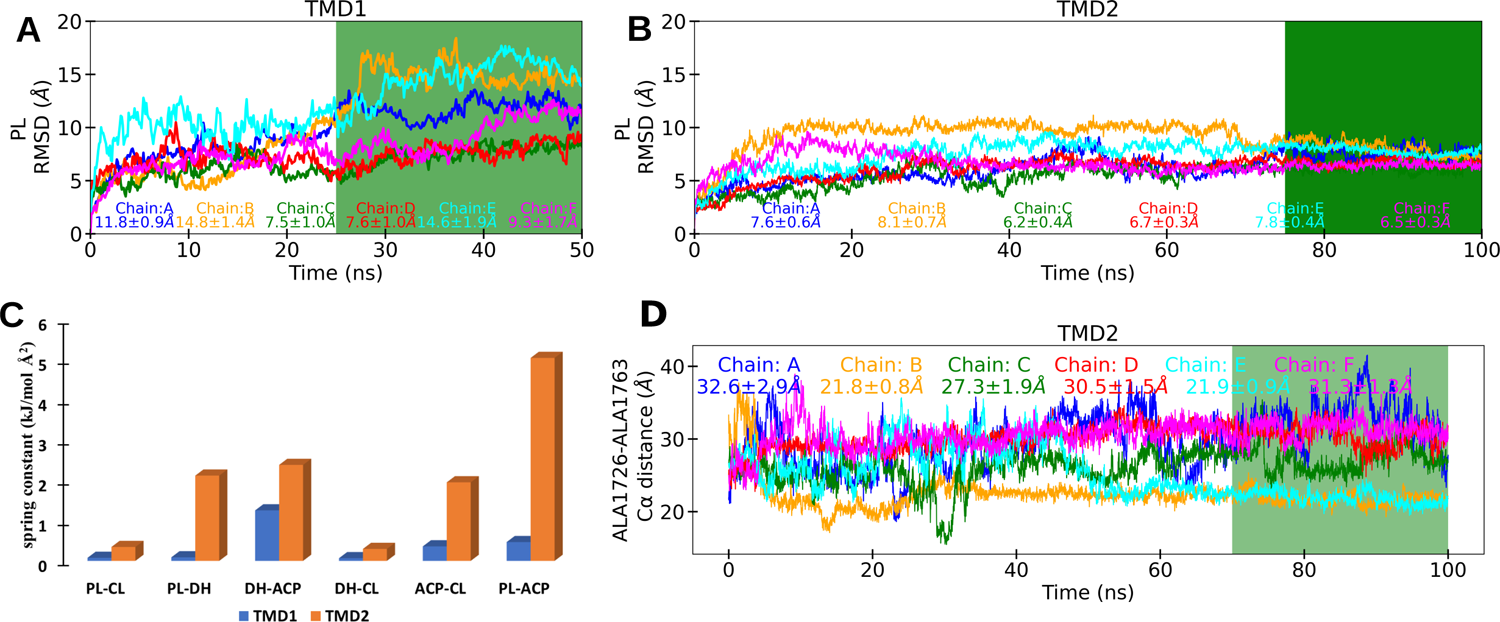
Comparison of PL dynamics through MD simulation and coarse-grained model. (A and B) RMSD of PL backbone atoms in each chain as a function of time, with mean and standard deviation in the last 25 ns (green region) included in the legend, indicating lower RMSD in traj2_100ns_ compared to traj1_50ns_ for all six chains. (C) Spring constants for domain-level interaction of PL with other domains obtained from the HENM model for both traj1_50ns_ and traj2_100ns_. (D) The distance between the ACP’s Cα of ALA1726 and Cα of PL ALA1763 in all six chains demonstrates the consistent interaction of PL with ACP in chain B.

In conclusion, homology modelling allowed us to determine the previously unknown structure of the mobile ACP in Mtb FAS-I, which was not observed in experimental cryo-EM reports at near-atomic resolution. Through classical MD simulation, we investigated the stability of the entire FAS-I complex with the modelled ACP structure stalled at DH, owing to lower occupancy. The stability of complex and ACP stalled at DH was aided by the unique conformation of PL in the reaction chamber, and its dynamics play a vital role in anchoring ACP near DH, which was unidentified in fungal or Mtb FAS-I complex. Based on that, we were able to observe ACP position in range to catalysis near DH in Mtb FAS-I i.e. distance between ACP and DH catalytic residue in between ∼18 Å to 22 Å, which is in good agreement with previous experimental results. Additionally, our computational analysis revealed correlated motion among the ER, MPT, and DH, and we observed that the adjacent ER was blocked due to ACP stalled at DH, presumably preventing other ACPs from interacting with other chain ERs. These findings contribute to a better understanding of the ACP shuttling mechanism and provide relevant insights into Mtb FAS-I complex for rational drug design.

## Acknowledgments

This research was supported in part by Science and Engineering Research Board (SERB) through the Start-up Research Grant SRG/2020/002186 and in part by Indian Institute of Technology Kanpur through the initiation grant. The computational support and the resources provided by PARAM Sanganak under the National Supercomputing Mission, Government of India at the Indian Institute of Technology Kanpur are gratefully acknowledged. A.K. also acknowledges financial support from Institute postdoctoral fellowship at Indian Institute of Technology Kanpur.

## 4 Author Contributions

H.H.K. designed the research. A.K carried out the MD simulations and analysed the results. M.S. carried out HENM analysis. A.K., M.S., and H.H.K co-wrote the manuscript.

## 6 STAR★Methods

### KEY RESOURCES TABLE

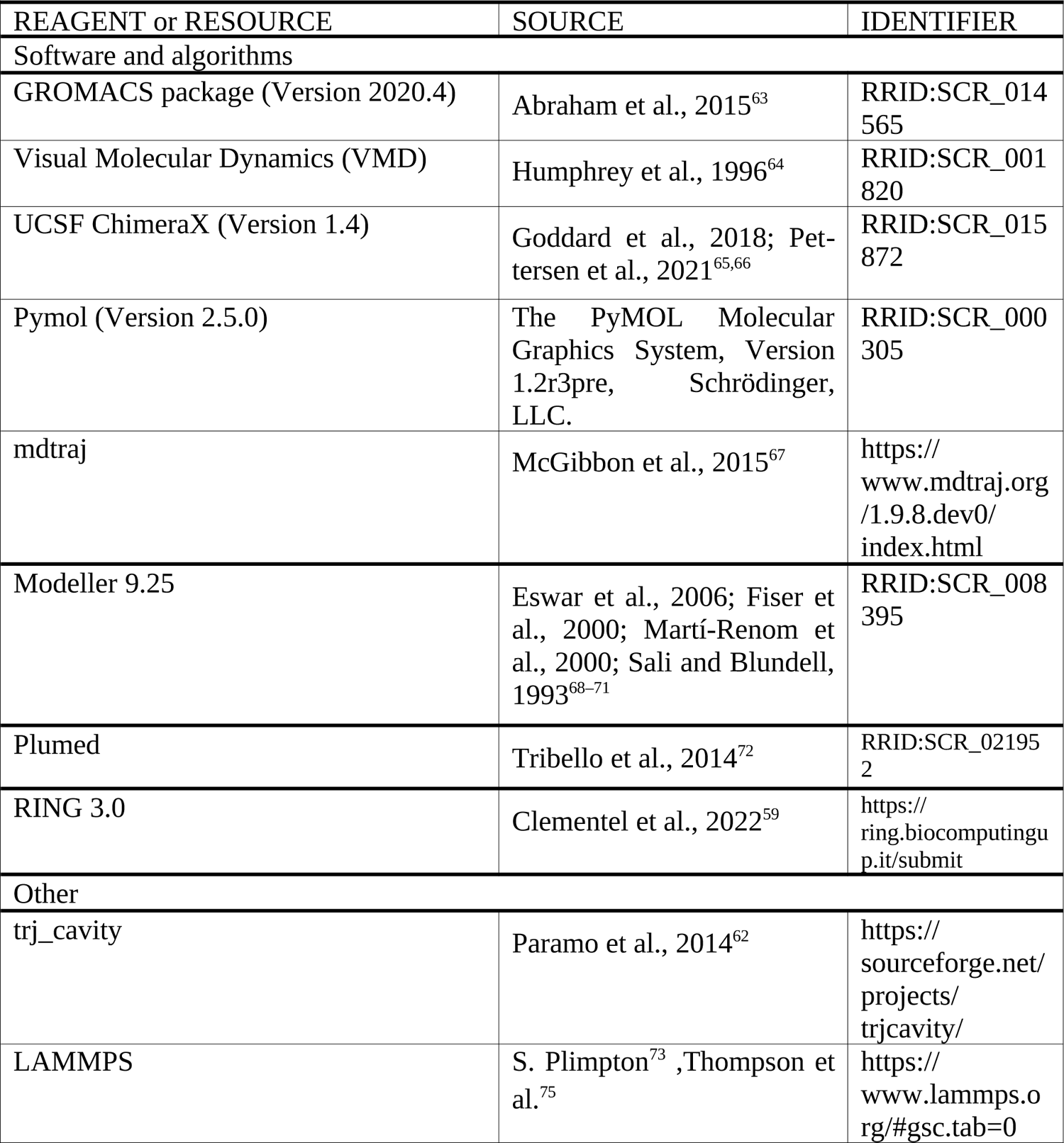

#### RESOURCE AVAILABILITY

##### Lead contact

Further information and requests for resources should be directed to and will be fulfilled by the lead contact Harshwardhan H. Katkar.

##### Materials availability

This study did not generate new unique reagents.

##### Data and code availability

Requests for the structural models and MD simulation trajectories should be directed to and will be fulfilled by the Lead contact, Harshwardhan H. Katkar.

This paper does not report original code.

Any additional information required to reanalyse the data reported in this paper is available from the Lead contact upon request.

## METHOD DETAILS

### 5.1 Homology modelling of Mtb-ACP; ACP stalled at KS

Mtb FAS-I structure at 3.3 Å (PDB ID: 6GJC)^50^ and its FASTA sequence (6GJC consists of six chains and each chain has 3096 residues) were retrieved from the protein data bank.^74^ The retrieved structure is missing ACP (1756-1924), linker region (PL:1710-1755, CL:1925-1942), and a small set of residues (2254-2262, 2303-2307) near the KR (numbering according to our model submitted with the manuscript) in all six chains. A BLASTP search did not reveal any known ACP structure in the protein data bank with a sequence homologous to Mtb ACP (also discussed in the Reviewers’ comments in Elad et al).^50^ The only exception is SC FAS-I, the structure of which was recently resolved at 3.1 Å using cryo-EM (PDB ID: 6TA1).^49^ This structure is inclusive of ACP stalled at the KS, with PL and CL regions still missing. Alignment of SC ACP and Mtb ACP using clustal 2.1 shows a 57% coverage and a 26.17% percentage identity (**Figure S5A**). The structure of a single chain of SC ACP was selected as a template to model the missing ACP structure of a single chain from Mtb FAS-I. The coordinates of this template were translated in order to overlay with the electron density of Mtb FAS-I before generating a homology model of the missing Mtb ACP structure, as described below.

Although the cryo-EM-based structure of Mtb FAS-I reported by Elad et al. at a resolution of 3.3 Å has missing regions inclusive of ACP, an unattributed relatively-weak electron density cloud has also been reported near the KS.^50^ The electron density map of Mtb FAS-I (EMD-0011) was retrieved from the electron microscopy data repository^76^ and analysed using the software package ChimeraX.^66^ Lowering the electron density threshold in ChimeraX to 0.0035, six prominent electron clouds of similar shape and size were observed near the KS: three in the upper dome and three in the lower dome (Figures 1B and 1C). In agreement with Elad et al., these were assigned as putative electron densities corresponding to ACP in the six chains of Mtb FAS-I.^50^

Upon aligning the structure of chain A and G of SC FAS-I from PDB ID 6TA1 with the structure of chain D of Mtb FAS-I using all atom of protein, a significant mismatch with the putative electron density of ACP was observed (Figures 1A and 1D**, SC ACP shown as gray tubes)**. To reduce this mismatch, the coordinates of all atoms belonging to ACP in the aligned structure were translated by 169.65 Å in the direction of [0.73, −0.66, 0.18]. The translated structure was used as a template for the homology modelling of Mtb ACP in a single chain. Ten models for Mtb ACP were generated using the segment modelling feature of Modeller 9.25^71^ **(Figure S5B)**. The model with the lowest dope score was selected **(Figure S5C)**. This model was further refined by remodelling for the loop region using loop modelling feature of Modeller 9.25 **(Figure S5C)**. PL and CL were subsequently modelled using the loop modelling feature of Modeller 9.25 (Figures 1A and 1D, ACP and PL shown as orange and purple tubes, respectively). This resulted into the complete, homology modelled structure of a single chain of Mtb FAS-I, which was then aligned with each of the six chains of 6GJC using all atoms of the rest of the domains to obtain the homology-modelled Mtb FAS-I hexamer (Figure 1E). Chains-IDs were assigned using the package PDB-tools.^77^

### 5.2 All-atom MD simulation of Mtb FAS-I; ACP stalled at KS

All-atom MD simulation of the homology modelled Mtb FAS-I hexamer was performed using GROMACS 2020.4.^63^ The hexamer structure was solvated using GROMACS utility *gmx solvate,* with a 30 Å x 30 Å x 30 Å periodic box. To neutralize the system, 732 Na^+^ ions were added to the system using GROMACS utility *gmx genion*. CHARMM36 force field^78^ with TIP3 water model^79^ was selected for the simulation. The system’s energy was minimized for 50000 steps using the steepest-descend algorithm. The resulting system was then equilibrated in two stages. In the first stage, the system was gradually heated from 0 K to 310 K using a linearly increasing temperature for 3 ns followed by 2 ns of equilibration at a constant volume and temperature (NVT ensemble) with a V-rescale thermostat and restraint (force constant 1000 kJ mol^-1^ nm^-2^) on all FAS-I atoms. In the second step, a total of 21 ns long constrained equilibration was performed at a pressure of 1 bar and temperature of 310 K with Parrinello-Rahman barostat and V-rescale thermostat (NPT ensemble), with constrain varying stepwise as follows: (a) 1 ns with restraint on all FAS-I atoms (force constant 1000 kJ mol^-1^ nm^-2^), (b) followed by 1 ns with restraint on all FAS-I atoms except hydrogen atoms (force constant 1000 kJ mol^-1^ nm^-2^), followed by (c) 5 ns with restraint on all FAS-I atoms except for atoms belonging to either ACP, PL or CL (force constant 1000 kJ mol^-1^ nm^-2^), followed by, (d) 14 ns with restraint on the backbone atoms of protein except ACP, PL and CL using varying force constant (**Figure S6B**). Figure 1F depicts the conformations of PL and ACP for chains D, E and F in the upper dome at the end of the 21 ns equilibration. The last frame of this simulation provided coordinates and velocities for the targeted MD simulations, TMD1 and TMD2, described in the following sections.

### 5.3 Targeted MD simulation of Mtb FAS-I; ACP stalled at DH (TMD1)

The structure of ACP stalled at DH in fungal FAS-I at a resolution of 5.80 Å was retrieved from the protein data bank (PDB ID: 6WC7).^80^ DH from the homology modelled structure of a single chain of Mtb FAS-I was aligned with that from a single chain of the fungal FAS-I using all atom of DH alone. This resulted into a single Mtb chain with ACP located near DH. The structure was aligned with each of the six chains in Mtb FAS-I retrieved from 6GJC using all atoms to generate a structure of Mtb FAS-I with ACP near DH in all six chains. This was used as a reference structure for the targeted MD simulation labelled TMD1 (Figure 1G), as described below.

The RMSD of Cα atoms in ACP, relative to the reference structure generated above, was chosen as the collective variable for performing a targeted MD simulation using GROMACS 2020.4 patched with PLUMED v2.3.^81^ The instantaneous RMSD was calculated using the RMSD command of the COLVAR module, and used to perform targeted MD simulation using the command MOVINGRESTRAINT of the BIAS module in PLUMED (**Figure S6A**). Initially, the RMSD was 5.5 nm. A time-dependent harmonic restraint was applied on the RMSD, with a force constant of 10000 kJ mol^-1^ nm^-2^ and a decreasing target value of RMSD over 5 ns as summarized in **Table S2.** At the end of TMD1, the RMSD was 0.91 nm. To prevent the movement of the rest of the domains (excluding ACP, PL and CL), we have applied harmonic restraint force using a force constant of 1000 kJ mol^-1^ nm^-2^ on all atoms of FAS-I. Figure 1H depicts the conformation of DH, ACP and PL in chains D, E, and F.

Following TMD1, the system was equilibrated for 5 ns using a constant volume and constant temperature of 310 K (NVT ensemble) with a V-rescale thermostat and a position restraint (force constant 1000 kJ mol^-1^ nm^-2^) on all backbone atoms of Mtb FAS-I, followed by 5 ns long equilibration at a constant pressure of 1 bar and temperature of 310 K with Parrinello-Rahman barostat and V-rescale thermostat (NPT ensemble), with the position restraints (force constant 1000 kJ mol^-1^ nm^-2^) on all backbone atoms. Subsequently, a 50 ns MD simulation trajectory was generated using the NPT ensemble in absence of any position restraints. Figure 1I depicts the conformation of DH, ACP and PL in chains D, E, F at end of 50 ns.

### 5.4 Targeted MD simulation of Mtb FAS-I; ACP with PL stalled at DH (TMD2)

In the 50 ns MD simulation following TMD1, ACP was found to interact with DH tenaciously in chain D, with a particular conformation of the PL stabilizing these interactions (**see section 3.1** for a detailed discussion). The structure of chain D at the end of the 50 ns simulation following TMD1 was aligned with each of the six chains in Mtb FAS-I retrieved from 6GJC using all atom to generate a structure of Mtb FAS-I with ACP near DH and PL in a particular conformation in all six chains. This was used as a reference structure (Figure 1J) for the targeted MD simulation labelled TMD2, as described below.

The RMSD of Cα atoms in ACP, relative to the reference structure generated above, was chosen as the collective variable for performing a targeted MD simulation using GROMACS 2020.4 patched with PLUMED v2.3.^72, 81^ The instantaneous RMSD was calculated using the RMSD command of the COLVAR module, and used to perform targeted MD simulation using the command MOVINGRESTRAINT of the BIAS module in PLUMED. Initially, the RMSD was 4.8 nm. A time-dependent harmonic restraint was applied on the RMSD, with a varying force constant and a decreasing target value of RMSD over 10 ns as summarized in **Table S2**. At the end of TMD1, the RMSD was 0.75 nm. To prevent the movement of the rest of the domains (excluding ACP, PL and CL) we have applied harmonic restraint force using a force constant of 1000 kJ mol^-1^ nm^-2^ on all atoms of FAS-I. Figure 1K depicts the conformations of PL, ACP and CL for chains D, E and F in the upper dome after TMD2.

Following TMD2, the system was equilibrated in two stages. The system was gradually heated from 0 K to 310 K using a linearly increasing temperature for 3 ns followed by 2 ns of equilibration at a constant volume and temperature (NVT ensemble) with a V-rescale thermostat and restraint (force constant 1000 kJ mol^-1^ nm^-2^) on all FAS-I atoms. In the second step, a total of 10-ns long constrained equilibration was performed at a pressure of 1 bar and temperature of 310 K with Parrinello-Rahman barostat and V-rescale thermostat (NPT ensemble), with constrain varying step wise as follows: (a) 1 ns with restraint on backbone FAS-I atoms (force constant 500 kJ mol ^-1^ nm^-2^), (b) followed by 10 ns with restraint on backbone atoms of protein except ACP, PL and CL using varying force constant (**Figure S6C**). Subsequently, a 100 ns MD simulation trajectory was generated using the NPT ensemble in the absence of any position restraint. Figure 1L depicts the conformations of PL, ACP and CL for chains D, E and F in the upper dome after 100 ns.

### 5.5 Fraction of native contacts

The fraction of native contacts is based on the definition provided by Best, Hummer and Eaton.^82^ Native pairs are defined as all the pairs of *i* and *j* heavy atoms belonging to residues *θ_i_* and *θ_j_* that are in contact, provided that along the primary sequence, |*θ_i_* − *θ_j_*| > 3 and the distance between *i* and *j, r_ij_,* is less than 4.5 Å. Based on this, the fraction of native contacts is defined as:

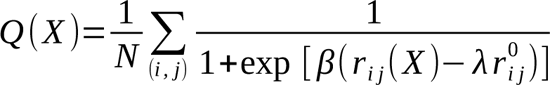

where, *N* is the number of pairs *i* and *j* that form native contacts, *r_ij_* (X) is the distance between *i* and *j* in configuration X, *r*° is the distance between *i* and *j* in the native state, *β* is the smoothing parameter with a value of 5 Å^-1^, and the factor *λ* accounts for fluctuations when contact is formed; it is taken as 1.8.

### 5.6 Heterogeneous elastic network model

In our work, we have employed a heterogeneous elastic network model (HENM) developed for characterization of fluctuations and quantify the correlations at domain-level structural fluctuations in the Mtb FAS-I complex. The coarse-grained (CG) force field was determined using an algorithm similar to that presented in Lyman et al.^83^ Normal mode analysis (NMA) was used to estimate the fluctuations of the CG sites.

Post-TMD simulations performed on Mtb FAS-I, traj1_50ns_ and traj2_100ns_, were analyzed using HENM to characterize the fluctuations at a multi-domain level. Mtb FAS-I complex consists of 7 domains in each homo-hexameric chain. The COM of all Cα in each domain has been assigned as one CG site (**Figure S7A**). PL and CL are considered as separate CG sites. We obtained the fluctuations in distances between all the CG pairs in CG MD mapped (CG-AA) trajectory for traj1_50ns_ and traj2_100ns_, <(Δr^2^_ij_)_CG-AA_>. The net harmonic interaction between all pairs of CG sites is given by:

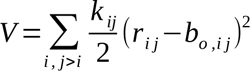

where, V is the total potential energy, k_ij_ is the spring constant for a CG pair, rij is the distance between a CG pair, and b_o_ij is the equilibrium distance between a CG pair. b_o_ij is estimated as the average distance between a CG pair in CG-AA trajectory.

We computed the normal modes for the CG model using the energy-minimized structure and calculate resulting fluctuations. We projected these fluctuations on energy minimized distance vectors *<*(Δ *r*^2^)*>*. The error between the fluctuations computed using normal mode analysis and target CG-AA mapped trajectory was minimized using Newton’s method and forward finite difference approximation, as

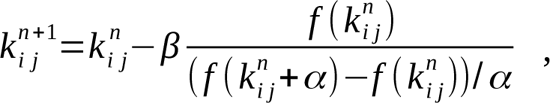

where,

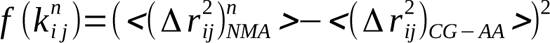

Here, n denotes the iteration, and parameter α and β controls the increment in spring constant and the size of the steps, respectively. We obtained an initial guess for the spring constants k_ij_ between the CG pairs using Boltzmann inversion. After minimizing the error, the obtained spring constants were used as the interaction potential between CG pairs to perform a Langevin dynamics simulation in LAMMPS molecular dynamics package^73, 75^ with a damp factor of 1 femtosecond. We verified that RMSF of the resulting 100 ns HENM-CG trajectory matches with the RMSF of CG-AA mapped trajectory (**Figure S7B**).

### 7.7 Simulation analysis

All simulations were visualized using Visual Molecular Dynamics (VMD)^64^ and PyMOL (The PyMOL Molecular Graphics System, Version 1.2r3pre, Schrödinger, LLC.). Other analysis tools were written with a combination of GROMACS tools and in house scripts, that utilised Python modules MDAnalysis and mdtraj. Figure and movie were prepared using chimeraX^66^ and VMD.

## 8 Supplemental information

Supplemental information (Document S1, S2 and Videos S1-S5) can be found online.

Document S1: Figures S1-S7, Table S1, S2 and Video legends S1-S5

Document S2: Pdb of the homology model Mtb FAS-I complex related to section 7.1

## References

1. Bloch, K., and Vance, D. (1977). Control Mechanisms in the Synthesis of Saturated Fatty Acids. Annual Review of Biochemistry 46, 263–298. 10.1146/annurev.bi.46.070177.001403.

2. Fernandes, N.D., and Kolattukudy, P.E. (1996). Cloning, sequencing and characterization of a fatty acid synthase-encoding gene from Mycobacterium tuberculosis var. bovis BCG. Gene 170, 95–99. 10.1016/0378-1119(95)00842-X.

3. Kikuchi, S., Rainwater, D.L., and Kolattukudy, P.E. (1992). Purification and characterization of an unusually large fatty acid synthase from Mycobacterium tuberculosis var. bovis BCG. Archives of Biochemistry and Biophysics 295, 318–326. 10.1016/0003-9861(92)90524-Z.

4. Wood, W.I., Peterson, D.O., and Bloch, K. (1978). Subunit structure of Mycobacterium smegmatis fatty acid synthetase. Evidence for identical multifunctional polypeptide chains. Journal of Biological Chemistry 253, 2650–2656. 10.1016/S0021-9258(17)40870-2.

5. Schweizer, E., and Hofmann, J. (2004). Microbial Type I Fatty Acid Synthases (FAS): Major Players in a Network of Cellular FAS Systems. Microbiol Mol Biol Rev 68, 501–517. 10.1128/MMBR.68.3.501-517.2004.

6. Sassetti, C.M., Boyd, D.H., and Rubin, E.J. (2001). Comprehensive identification of conditionally essential genes in mycobacteria. Proc. Natl. Acad. Sci. U.S.A. 98, 12712–12717. 10.1073/pnas.231275498.

7. Zimhony, O., Vilchèze, C., and Jacobs, W.R. (2004). Characterization of *Mycobacterium smegmatis* Expressing the *Mycobacterium tuberculosis* Fatty Acid Synthase I (*fas1*) Gene. J Bacteriol 186, 4051–4055. 10.1128/JB.186.13.4051-4055.2004.

8. Takayama, K., Wang, C., and Besra, G.S. (2005). Pathway to Synthesis and Processing of Mycolic Acids in *Mycobacterium tuberculosis*. Clin Microbiol Rev 18, 81–101. 10.1128/CMR.18.1.81-101.2005.

9. Brennan, P.J., and Nikaido, H. (1995). THE ENVELOPE OF MYCOBACTERIA. Annu. Rev. Biochem. 64, 29–63. 10.1146/annurev.bi.64.070195.000333.

10. Lynen, F., Engeser, H., Foerster, E.-C., Fox, J.L., Hess, S., Kresze, G.-B., Schmitt, T., Schreckenbach, T., Siess, E., Wieland, F., et al. (1980). On the Structure of Fatty Acid Synthetase of Yeast. European Journal of Biochemistry 112, 431–442. 10.1111/j.1432-1033.1980.tb06105.x.

11. Stoops, J.K., Wakil, S.J., Uberbacher, E.C., and Bunick, G.J. (1987). Small-angle neutron-scattering and electron microscope studies of the chicken liver fatty acid synthase. J Biol Chem 262, 10246–10251.

12. Jenni, S., Leibundgut, M., Maier, T., and Ban, N. (2006). Architecture of a fungal fatty acid synthase at 5 A resolution. Science 311, 1263–1267. 10.1126/science.1123251.

13. Ciccarelli, L., Connell, S.R., Enderle, M., Mills, D.J., Vonck, J., and Grininger, M. (2013). Structure and Conformational Variability of the Mycobacterium tuberculosis Fatty Acid Synthase Multienzyme Complex. Structure 21, 1251–1257. 10.1016/j.str.2013.04.023.

14. Boehringer, D., Ban, N., and Leibundgut, M. (2013). 7.5-Å Cryo-EM Structure of the Mycobacterial Fatty Acid Synthase. Journal of Molecular Biology 425, 841–849. 10.1016/j.jmb.2012.12.021.

15. Stoops, J.K., Awad, E.S., Arslanian, M.J., Gunsberg, S., Wakil, S.J., and Oliver, R.M. (1978). Studies on the yeast fatty acid synthetase. Subunit composition and structural organization of a large multifunctional enzyme complex. Journal of Biological Chemistry 253, 4464–4475. 10.1016/S0021-9258(17)34743-9.

16. Stoops, J.K., and Wakil, S.J. (1980). Yeast fatty acid synthetase: structure-function relationship and nature of the beta-ketoacyl synthetase site. Proc Natl Acad Sci U S A 77, 4544–4548. 10.1073/pnas.77.8.4544.

17. Lou, J.W., and Mazhab-Jafari, M.T. (2020). Steric occlusion regulates proximal interactions of acyl carrier protein domain in fungal fatty acid synthase. Commun Biol 3, 274. 10.1038/s42003-020-0997-y.

18. Maier, T., Leibundgut, M., Boehringer, D., and Ban, N. (2010). Structure and function of eukaryotic fatty acid synthases. Quarterly Reviews of Biophysics 43, 373–422. 10.1017/S0033583510000156.

19. Leibundgut, M., Maier, T., Jenni, S., and Ban, N. (2008). The multienzyme architecture of eukaryotic fatty acid synthases. Curr Opin Struct Biol 18, 714–725. 10.1016/j.sbi.2008.09.008.

20. Wakil, S.J. (1989). Fatty acid synthase, a proficient multifunctional enzyme. Biochemistry 28, 4523–4530. 10.1021/bi00437a001.

21. Fernandes, N.D., and Kolattukudy, P.E. (1996). Cloning, sequencing and characterization of a fatty acid synthase-encoding gene from Mycobacterium tuberculosis var. bovis BCG. Gene 170, 95–99. 10.1016/0378-1119(95)00842-x.

22. Byers, D.M., and Gong, H. (2007). Acyl carrier protein: structure–function relationships in a conserved multifunctional protein family. Biochem. Cell Biol. 85, 649–662. 10.1139/O07-109.

23. Majerus, P.W., Alberts, A.W., and Vagelos, P.R. (1965). ACYL CARRIER PROTEIN, IV. THE IDENTIFICATION OF 4′-PHOSPHOPANTETHEINE AS THE PROSTHETIC GROUP OF THE ACYL CARRIER PROTEIN. Proc Natl Acad Sci U S A 53, 410–417.

24. Roncari, D.A.K., Bradshaw, R.A., and Vagelos, P.R. (1972). Acyl Carrier Protein: XIX. AMINO ACID SEQUENCE OF LIVER FATTY ACID SYNTHETASE PEPTIDES CONTAINING 4′-PHOSPHOPANTETHEINE. Journal of Biological Chemistry 247, 6234–6242. 10.1016/S0021-9258(19)44787-X.

25. Willecke, K., Ritter, E., and Lynen, F. (1969). Isolation of an Acyl Carrier Protein Component from the Multienzyme Complex of Yeast Fatty Acid Synthetase. European Journal of Biochemistry 8, 503–509. 10.1111/j.1432-1033.1969.tb00555.x.

26. Gipson, P., Mills, D.J., Wouts, R., Grininger, M., Vonck, J., and Kühlbrandt, W. (2010). Direct structural insight into the substrate-shuttling mechanism of yeast fatty acid synthase by electron cryomicroscopy. Proc. Natl. Acad. Sci. U.S.A. 107, 9164–9169. 10.1073/pnas.0913547107.

27. Jenni, S., Leibundgut, M., Boehringer, D., Frick, C., Mikolásek, B., and Ban, N. (2007). Structure of Fungal Fatty Acid Synthase and Implications for Iterative Substrate Shuttling. Science 316, 254–261. 10.1126/science.1138248.

28. Lomakin, I.B., Xiong, Y., and Steitz, T.A. (2007). The Crystal Structure of Yeast Fatty Acid Synthase, a Cellular Machine with Eight Active Sites Working Together. Cell 129, 319–332. 10.1016/j.cell.2007.03.013.

29. Fischer, M., Joppe, M., Mulinacci, B., Vollrath, R., Konstantinidis, K., Kötter, P., Ciccarelli, L., Vonck, J., Oesterhelt, D., and Grininger, M. (2020). Analysis of the co-translational assembly of the fungal fatty acid synthase (FAS). Sci Rep 10, 895. 10.1038/s41598-020-57418-8.

30. Kastritis, P.L., O’Reilly, F.J., Bock, T., Li, Y., Rogon, M.Z., Buczak, K., Romanov, N., Betts, M.J., Bui, K.H., Hagen, W.J., et al. (2017). Capturing protein communities by structural proteomics in a thermophilic eukaryote. Mol Syst Biol 13, 936. 10.15252/msb.20167412.

31. Leibundgut, M., Jenni, S., Frick, C., and Ban, N. (2007). Structural Basis for Substrate Delivery by Acyl Carrier Protein in the Yeast Fatty Acid Synthase. Science 316, 288–290. 10.1126/science.1138249.

32. Fischer, M., Rhinow, D., Zhu, Z., Mills, D.J., Zhao, Z.K., Vonck, J., and Grininger, M. (2015). Cryo-EM structure of fatty acid synthase (FAS) from Rhodosporidium toruloides provides insights into the evolutionary development of fungal FAS. Protein Science 24, 987–995. 10.1002/pro.2678.

33. Anselmi, C., Grininger, M., Gipson, P., and Faraldo-Gómez, J.D. (2010). Mechanism of Substrate Shuttling by the Acyl-Carrier Protein within the Fatty Acid Mega-Synthase. J. Am. Chem. Soc. 132, 12357–12364. 10.1021/ja103354w.

34. Pugh, E.L., and Wakil, S.J. (1965). Studies on the Mechanism of Fatty Acid Synthesis: XIV. THE PROSTHETIC GROUP OF ACYL CARRIER PROTEIN AND THE MODE OF ITS ATTACHMENT TO THE PROTEIN. Journal of Biological Chemistry 240, 4727–4733. 10.1016/S0021-9258(18)97016-X.

35. Roujeinikova, A., Simon, W.J., Gilroy, J., Rice, D.W., Rafferty, J.B., and Slabas, A.R. (2007). Structural Studies of Fatty Acyl-(Acyl Carrier Protein) Thioesters Reveal a Hydrophobic Binding Cavity that Can Expand to Fit Longer Substrates. Journal of Molecular Biology 365, 135–145. 10.1016/j.jmb.2006.09.049.

36. Tsai, S.-C. (Sheryl), and Ames, B.D. (2009). STRUCTURAL ENZYMOLOGY OF POLYKETIDE SYNTHASES. Methods Enzymol 459, 17–47. 10.1016/S0076-6879(09)04602-3.

37. Zornetzer, G.A., Fox, B.G., and Markley, J.L. (2006). Solution structures of spinach acyl carrier protein with decanoate and stearate. Biochemistry 45, 5217–5227. 10.1021/bi052062d.

38. Lou, J.W., Iyer, K.R., Hasan, S.M.N., Cowen, L.E., and Mazhab-Jafari, M.T. (2019). Electron cryomicroscopy observation of acyl carrier protein translocation in type I fungal fatty acid synthase. Sci Rep 9, 1–8. 10.1038/s41598-019-49261-3.

39. Qiu, S., Liu, S., Zaoti, Z.F., Wang, X., and Cai, G. (2019). Modulation of fatty acid synthase by ATR checkpoint kinase Rad3. J Mol Cell Biol 11, 1098–1100. 10.1093/jmcb/mjz096.

40. Fichtlscherer, F., Wellein, C., Mittag, M., and Schweizer, E. (2000). A novel function of yeast fatty acid synthase. Subunit alpha is capable of self-pantetheinylation. Eur J Biochem 267, 2666–2671. 10.1046/j.1432-1327.2000.01282.x.

41. Mohamed, A.H., Chirala, S.S., Mody, N.H., Huang, W.Y., and Wakil, S.J. (1988). Primary structure of the multifunctional alpha subunit protein of yeast fatty acid synthase derived from FAS2 gene sequence. J Biol Chem 263, 12315–12325.

42. Schreckenbach, T., Wobser, H., and Lynen, F. (1977). The palmityl binding sites of fatty acid synthetase from yeast. Eur J Biochem 80, 13–23. 10.1111/j.1432-1033.1977.tb11850.x.

43. Schuster, H., Rautenstrauss, B., Mittag, M., Stratmann, D., and Schweizer, E. (1995). Substrate and product binding sites of yeast fatty acid synthase. Stoichiometry and binding kinetics of wild-type and in vitro mutated enzymes. Eur J Biochem 228, 417–424.

44. Parris, K.D., Lin, L., Tam, A., Mathew, R., Hixon, J., Stahl, M., Fritz, C.C., Seehra, J., and Somers, W.S. (2000). Crystal structures of substrate binding to Bacillus subtilis holo-(acyl carrier protein) synthase reveal a novel trimeric arrangement of molecules resulting in three active sites. Structure 8, 883–895. 10.1016/S0969-2126(00)00178-7.

45. Rafi, S., Novichenok, P., Kolappan, S., Stratton, C.F., Rawat, R., Kisker, C., Simmerling, C., and Tonge, P.J. (2006). Structure of acyl carrier protein bound to FabI, the FASII enoyl reductase from Escherichia coli. J Biol Chem 281, 39285–39293. 10.1074/jbc.M608758200.

46. Zhang, Y.-M., Wu, B., Zheng, J., and Rock, C.O. (2003). Key residues responsible for acyl carrier protein and beta-ketoacyl-acyl carrier protein reductase (FabG) interaction. J Biol Chem 278, 52935–52943. 10.1074/jbc.M309874200.

47. Zhang, Y.M., Rao, M.S., Heath, R.J., Price, A.C., Olson, A.J., Rock, C.O., and White, S.W. (2001). Identification and analysis of the acyl carrier protein (ACP) docking site on beta-ketoacyl-ACP synthase III. J Biol Chem 276, 8231–8238. 10.1074/jbc.M008042200.

48. Medina, F.E., Neves, R.P.P., Ramos, M.J., and Fernandes, P.A. (2018). QM/MM Study of the Reaction Mechanism of the Dehydratase Domain from Mammalian Fatty Acid Synthase. ACS Catal. 8, 10267–10278. 10.1021/acscatal.8b02616.

49. Joppe, M., D’Imprima, E., Salustros, N., Paithankar, K.S., Vonck, J., Grininger, M., and Kühlbrandt, W. (2020). The resolution revolution in cryoEM requires high-quality sample preparation: a rapid pipeline to a high-resolution map of yeast fatty acid synthase. IUCrJ 7, 220–227. 10.1107/S2052252519017366.

50. Elad, N., Baron, S., Peleg, Y., Albeck, S., Grunwald, J., Raviv, G., Shakked, Z., Zimhony, O., and Diskin, R. (2018). Structure of Type-I Mycobacterium tuberculosis fatty acid synthase at 3.3 Å resolution. Nat Commun 9, 3886. 10.1038/s41467-018-06440-6.

51. Masoudi, A., Raetz, C.R.H., Zhou, P., and Pemble, C.W. (2014). Chasing Acyl-Carrier-Protein Through a Catalytic Cycle of Lipid A Production. Nature 505, 422–426. 10.1038/nature12679.

52. Wong, H.C., Liu, G., Zhang, Y.-M., Rock, C.O., and Zheng, J. (2002). The Solution Structure of Acyl Carrier Protein from Mycobacterium tuberculosis*. Journal of Biological Chemistry 277, 15874–15880. 10.1074/jbc.M112300200.

53. Xu, G.Y., Tam, A., Lin, L., Hixon, J., Fritz, C.C., and Powers, R. (2001). Solution structure of B. subtilis acyl carrier protein. Structure 9, 277–287. 10.1016/s0969-2126(01)00586-x.

54. Płoskoń, E., Arthur, C.J., Kanari, A.L.P., Wattana-amorn, P., Williams, C., Crosby, J., Simpson, T.J., Willis, C.L., and Crump, M.P. (2010). Recognition of Intermediate Functionality by Acyl Carrier Protein over a Complete Cycle of Fatty Acid Biosynthesis. Chemistry & Biology 17, 776–785. 10.1016/j.chembiol.2010.05.024.

55. White, S.W., Zheng, J., Zhang, Y.-M., and Rock, null (2005). The structural biology of type II fatty acid biosynthesis. Annu Rev Biochem 74, 791–831. 10.1146/annurev.biochem.74.082803.133524.

56. Colizzi, F., Recanatini, M., and Cavalli, A. (2008). Mechanical Features of Plasmodium falciparum Acyl Carrier Protein in the Delivery of Substrates. J. Chem. Inf. Model. 48, 2289– 2293. 10.1021/ci800297v.

57. Sharma, A.K., Sharma, S.K., Surolia, A., Surolia, N., and Sarma, S.P. (2006). Solution Structures of Conformationally Equilibrium Forms of Holo-Acyl Carrier Protein (PfACP) from Plasmodium falciparum Provides Insight into the Mechanism of Activation of ACPs,. Biochemistry 45, 6904–6916. 10.1021/bi060368u.

58. Chan, D.I., Stockner, T., Tieleman, D.P., and Vogel, H.J. (2008). Molecular Dynamics Simulations of the Apo-, Holo-, and Acyl-forms of Escherichia coli Acyl Carrier Protein. J Biol Chem 283, 33620–33629. 10.1074/jbc.M805323200.

59. Clementel, D., Del Conte, A., Monzon, A.M., Camagni, G.F., Minervini, G., Piovesan, D., and Tosatto, S.C.E. (2022). RING 3.0: fast generation of probabilistic residue interaction networks from structural ensembles. Nucleic Acids Research 50, W651–W656. 10.1093/nar/gkac365.

60. Flaman, A.S., Chen, J.M., Van Iderstine, S.C., and Byers, D.M. (2001). Site-directed mutagenesis of acyl carrier protein (ACP) reveals amino acid residues involved in ACP structure and acyl-ACP synthetase activity. J Biol Chem 276, 35934–35939. 10.1074/jbc.M101849200.

61. Leesong, M., Henderson, B.S., Gillig, J.R., Schwab, J.M., and Smith, J.L. (1996). Structure of a dehydratase–isomerase from the bacterial pathway for biosynthesis of unsaturated fatty acids: two catalytic activities in one active site. Structure 4, 253–264. 10.1016/S0969-2126(96)00030-5.

62. Paramo, T., East, A., Garzón, D., Ulmschneider, M.B., and Bond, P.J. (2014). Efficient Characterization of Protein Cavities within Molecular Simulation Trajectories: trj_cavity. J. Chem. Theory Comput. 10, 2151–2164. 10.1021/ct401098b.

63. Abraham, M.J., Murtola, T., Schulz, R., Páll, S., Smith, J.C., Hess, B., and Lindahl, E. (2015). GROMACS: High performance molecular simulations through multi-level parallelism from laptops to supercomputers. SoftwareX 1–2, 19–25. 10.1016/j.softx.2015.06.001.

64. Humphrey, W., Dalke, A., and Schulten, K. (1996). VMD: visual molecular dynamics. J Mol Graph 14, 33–38, 27–28. 10.1016/0263-7855(96)00018-5.

65. Goddard, T.D., Huang, C.C., Meng, E.C., Pettersen, E.F., Couch, G.S., Morris, J.H., and Ferrin, T.E. (2018). UCSF ChimeraX: Meeting modern challenges in visualization and analysis. Protein Sci 27, 14–25. 10.1002/pro.3235.

66. Pettersen, E.F., Goddard, T.D., Huang, C.C., Meng, E.C., Couch, G.S., Croll, T.I., Morris, J.H., and Ferrin, T.E. (2021). UCSF ChimeraX: Structure visualization for researchers, educators, and developers. Protein Sci 30, 70–82. 10.1002/pro.3943.

67. McGibbon, R.T., Beauchamp, K.A., Harrigan, M.P., Klein, C., Swails, J.M., Hernández, C.X., Schwantes, C.R., Wang, L.-P., Lane, T.J., and Pande, V.S. (2015). MDTraj: A Modern Open Library for the Analysis of Molecular Dynamics Trajectories. Biophys J 109, 1528–1532. 10.1016/j.bpj.2015.08.015.

68. Eswar, N., Webb, B., Marti-Renom, M.A., Madhusudhan, M.S., Eramian, D., Shen, M.-Y., Pieper, U., and Sali, A. (2006). Comparative protein structure modeling using Modeller. Curr Protoc Bioinformatics Chapter 5, Unit-5.6. 10.1002/0471250953.bi0506s15.

69. Fiser, A., Do, R.K.G., and Šali, A. (2000). Modeling of loops in protein structures. Protein Science 9, 1753–1773. 10.1110/ps.9.9.1753.

70. Martí-Renom, M.A., Stuart, A.C., Fiser, A., Sánchez, R., Melo, F., and Sali, A. (2000). Comparative protein structure modeling of genes and genomes. Annu Rev Biophys Biomol Struct 29, 291–325. 10.1146/annurev.biophys.29.1.291.

71. Sali, A., and Blundell, T.L. (1993). Comparative protein modelling by satisfaction of spatial restraints. J Mol Biol 234, 779–815. 10.1006/jmbi.1993.1626.

72. Tribello, G.A., Bonomi, M., Branduardi, D., Camilloni, C., and Bussi, G. (2014). PLUMED 2: New feathers for an old bird. Computer Physics Communications 185, 604–613. 10.1016/j.cpc.2013.09.018.

73. Plimpton, S. (1995). Fast Parallel Algorithms for Short-Range Molecular Dynamics. Journal of Computational Physics 117, 1–19. 10.1006/jcph.1995.1039.

74. Berman, H.M., Westbrook, J., Feng, Z., Gilliland, G., Bhat, T.N., Weissig, H., Shindyalov, I.N., and Bourne, P.E. (2000). The Protein Data Bank. Nucleic Acids Research 28, 235–242. 10.1093/nar/28.1.235.

75. Thompson, A.P., Aktulga, H.M., Berger, R., Bolintineanu, D.S., Brown, W.M., Crozier, P.S., in’t Veld, P.J., Kohlmeyer, A., Moore, S.G., Nguyen, T.D., et al. (2022). LAMMPS - a flexible simulation tool for particle-based materials modeling at the atomic, meso, and continuum scales. Computer Physics Communications 271, 108171. 10.1016/j.cpc.2021.108171.

76. EMDB Electron Microscopy Data Bank. Electron Microscopy Data Bank. https://www.ebi.ac.uk/emdb/EMD-0011.

77. Rodrigues, J.P.G.L.M., Teixeira, J.M.C., Trellet, M., and Bonvin, A.M.J.J. (2018). pdb-tools: a swiss army knife for molecular structures. F1000Res 7, 1961. 10.12688/f1000research.17456.1.

78. Huang, J., and MacKerell, A.D. (2013). CHARMM36 all-atom additive protein force field: validation based on comparison to NMR data. J Comput Chem 34, 2135–2145. 10.1002/jcc.23354.

79. Jorgensen, W.L., Chandrasekhar, J., Madura, J.D., Impey, R.W., and Klein, M.L. (1983). Comparison of simple potential functions for simulating liquid water. J. Chem. Phys. 79, 926–935. 10.1063/1.445869.

80. Lou, J.W., and Mazhab-Jafari, M.T. (2020). Steric occlusion regulates proximal interactions of acyl carrier protein domain in fungal fatty acid synthase. Commun Biol 3, 1–9. 10.1038/s42003-020-0997-y.

81. Bonomi, M., Branduardi, D., Bussi, G., Camilloni, C., Provasi, D., Raiteri, P., Donadio, D., Marinelli, F., Pietrucci, F., Broglia, R.A., et al. (2009). PLUMED: A portable plugin for free-energy calculations with molecular dynamics. Computer Physics Communications 180, 1961– 1972. 10.1016/j.cpc.2009.05.011.

82. Best, R.B., Hummer, G., and Eaton, W.A. (2013). Native contacts determine protein folding mechanisms in atomistic simulations. Proceedings of the National Academy of Sciences 110, 17874–17879. 10.1073/pnas.1311599110.

83. Lyman, E., Pfaendtner, J., and Voth, G.A. (2008). Systematic Multiscale Parameterization of Heterogeneous Elastic Network Models of Proteins. Biophysical Journal 95, 4183–4192. 10.1529/biophysj.108.139733.

