## Supplementary material for "Peripheral linker mediates ACP’s recognition of DH and stabilizes Mycobacterium tuberculosis FAS-I": Document S1

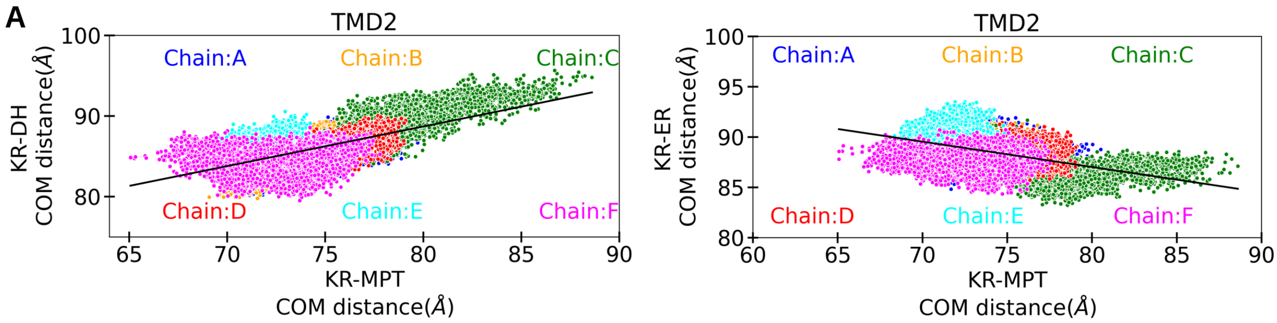


**Figure S1. Coordinated interplay between MPT, DH, and ER (related to Figure 3).** (A) Positive correlation between MPT and DH and (B) Negative correlation between MPT and ER in traj2$_{100ns}$. The lines shows the linear regression.

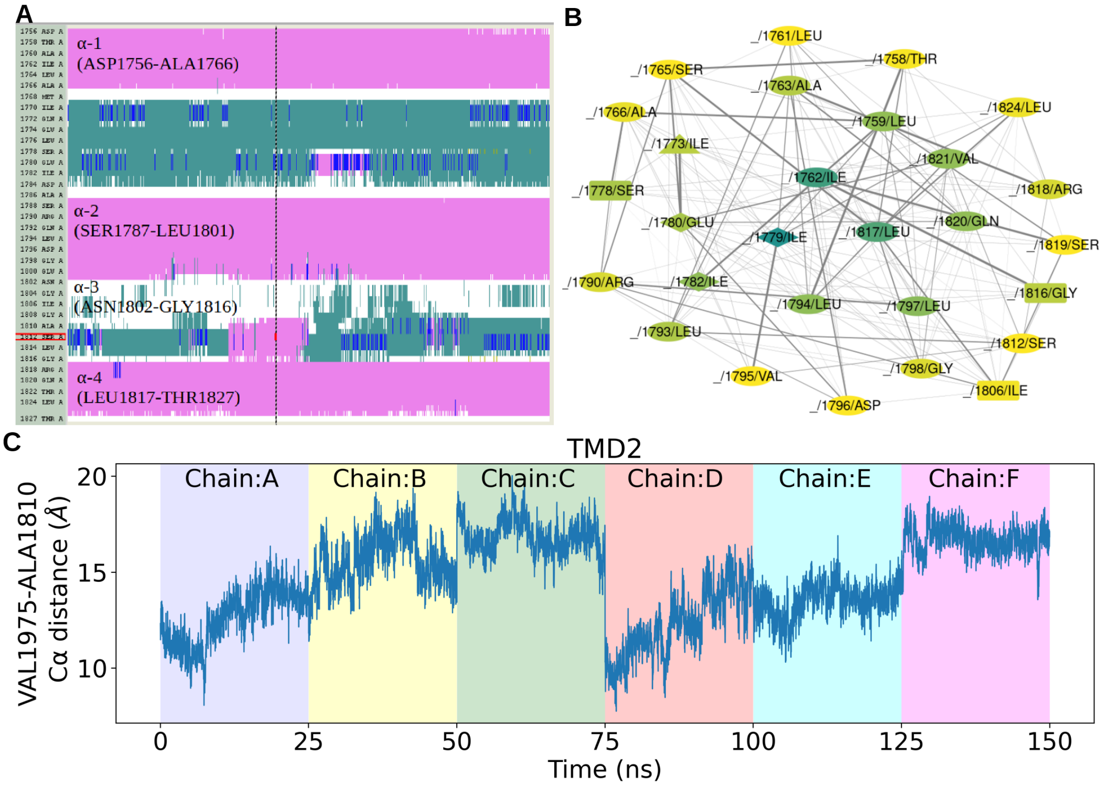


**Figure S2. Secondary structure analysis and interaction for catalytic core of Mtb ACP for last 25 ns of traj2$_{100ns}$ for each chain (related to Figure 4).** (A) Three solid pink region shows helices α-1, α-2, α-4 adopt alpha helix conformation; amino acid range from ASN1802 to GLY1816 i.e., α-3 *w*hich adopt alpha helix (pink color) and 3-10 helix (blue color) and turn (green color) conformation, white color represent coil structure. (B) Network analysis for the residue level interaction in ACP catalytic core. (C) Evolution of distance between Cα atoms of VAL1975 and ALA1810 of ACP catalytic core for all six chains for last 25 ns. Minimum distance and maximum distance between Cα atom of these two residues is reported to be 20 Å that is in agreement of previous MD simulation of PL-ACP.[1]

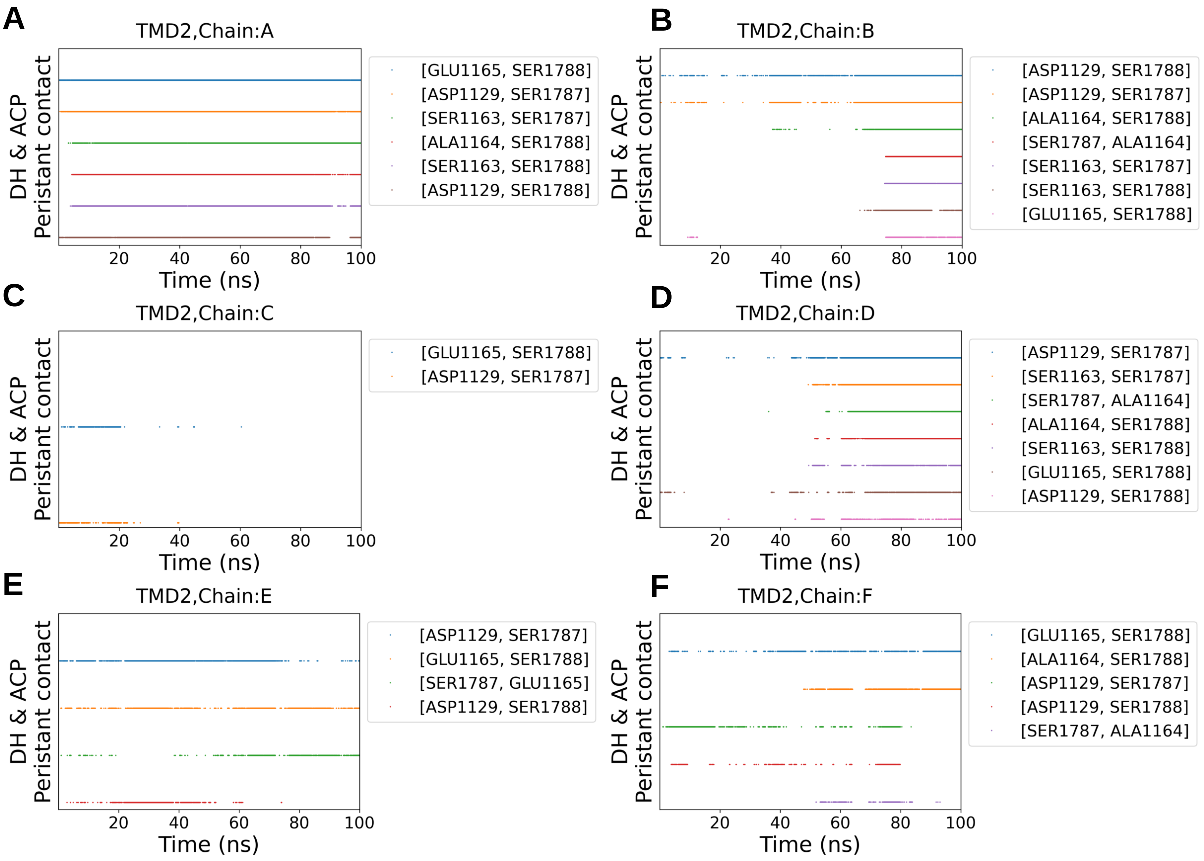


**Figure S3. Key surface residues involve persistent DH-ACP interaction in traj2$_{100ns}$ (related to Figure 5).** (A-F) Persistent contact in all six chains between ACP catalytic core and DH surface residues in Mtb FAS-I complex in traj2$_{100ns}$.

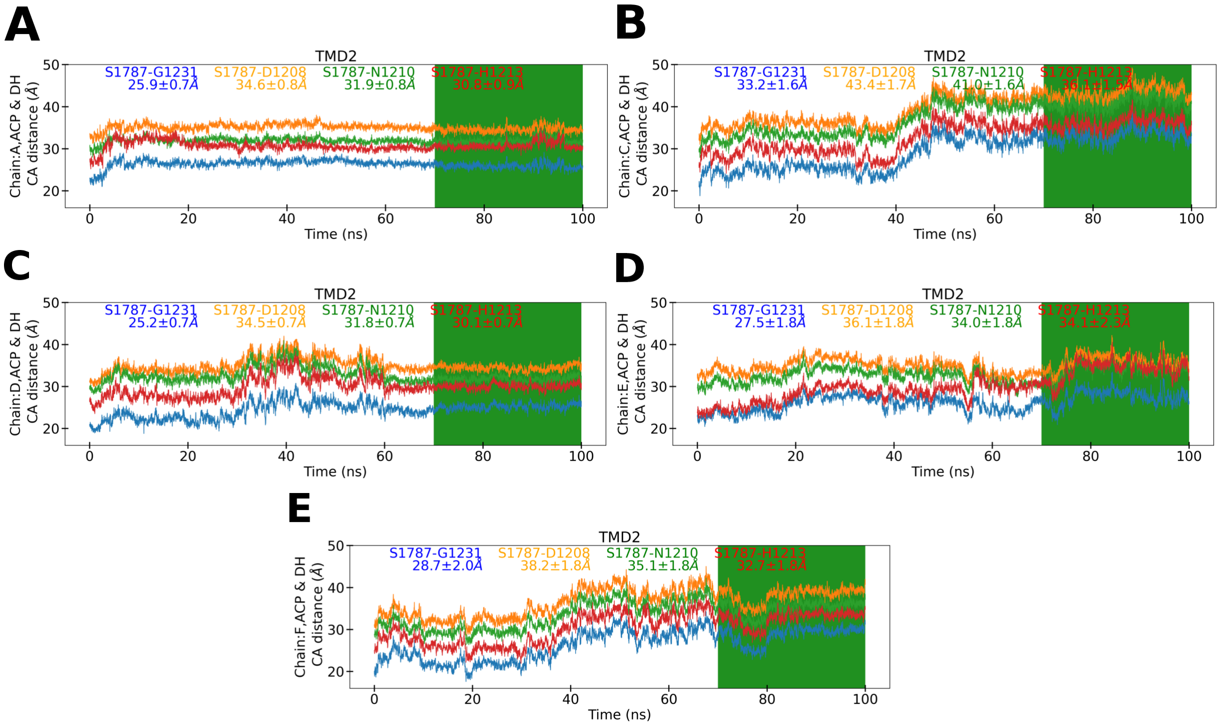


**Figure S4. Distances between catalytic residues of DH and ACP SER1787 in traj2$_{100ns}$ (related to Figure 5 ).** (A-E) Distance of DH residues (CG of ASP1208, NG1 of HIS1213, CA of ASN1210, CA of GLY1231) with Cβ of SER1787 of ACP.

**A**

```
CLUSTAL 2.1 multiple sequence alignment


pdb_Y          AAEIADEPVKASLLLHVLVAHKLKKSLDSIPMSKTIKDLVGGKSTVQNEILGDLGKEFGT
pdb_M          ------------DATLALIALSAKMRIDQIEELDSIESITDGASSRRNQLLVDLGSELNL
                               .*:* . *  :*.*   .:*:.:..* *:  :*::* ***.*:.

pdb_Y          ----TPEKPEETPLEELAETFQDTFSGALGKQSSSLLSRLISSKMPGGFTITVARKYLQT
pdb_M          GAIDGAAESDLAGLRSQVTKLARTYKPYGPVLSDAINDQLRTVLGPSGKRPGAIAERVKK
                    . :.: : *.. . .:  *:.      *.:: .:* :   *.*    .  : ::.

pdb_Y          RWGLPSGRQDGVLLVALSNEPAAR-----------LGSEADAKAFLDSMAQKYASIVGVD
pdb_M          TWELGEGWAKHVTVEVALGTREGSSVRGGAMGHLHEGALADAASVDKVIDAAVASVAARQ
                * * .*  . * : .  .   .             *: *** :. . :    **:.. :

pdb_Y          L
pdb_M          G
```

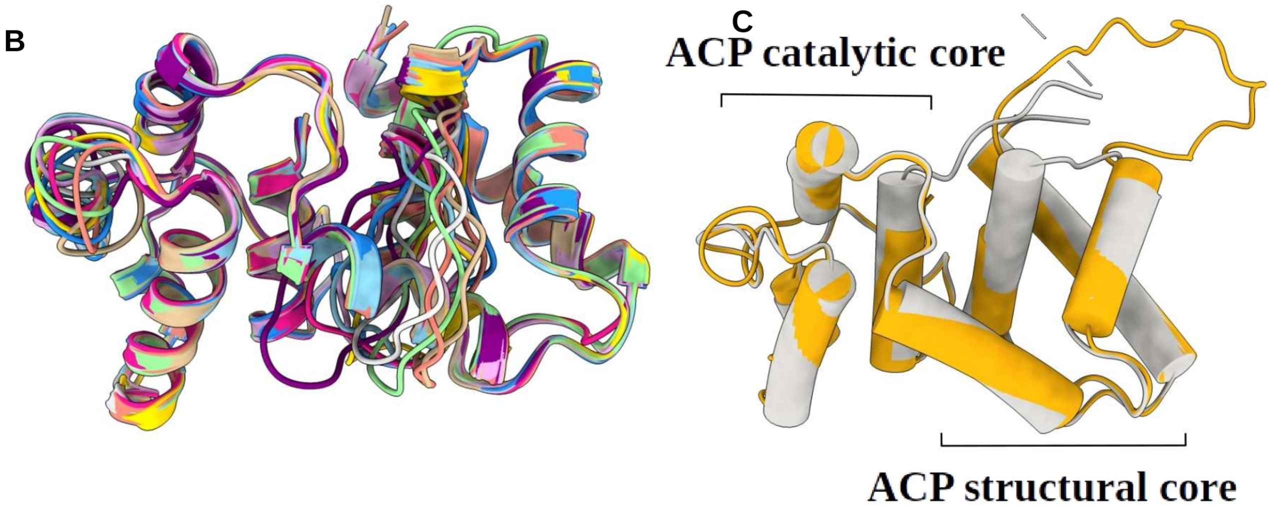


**Figure S5. Prediction of Mtb FAS-I ACP (related to Figure 4 ).** (A) Homology modelling of Mtb ACP; ACP stalled at KS: showing ten superimposed model for Mtb ACP, generated using Modeller 9.25 software. (B) Comparison between template 6TA1 and final model after refinement. (C) Catalytic and structural core helices of ACP of fungal FAS-I and modelled ACP of Mtb.

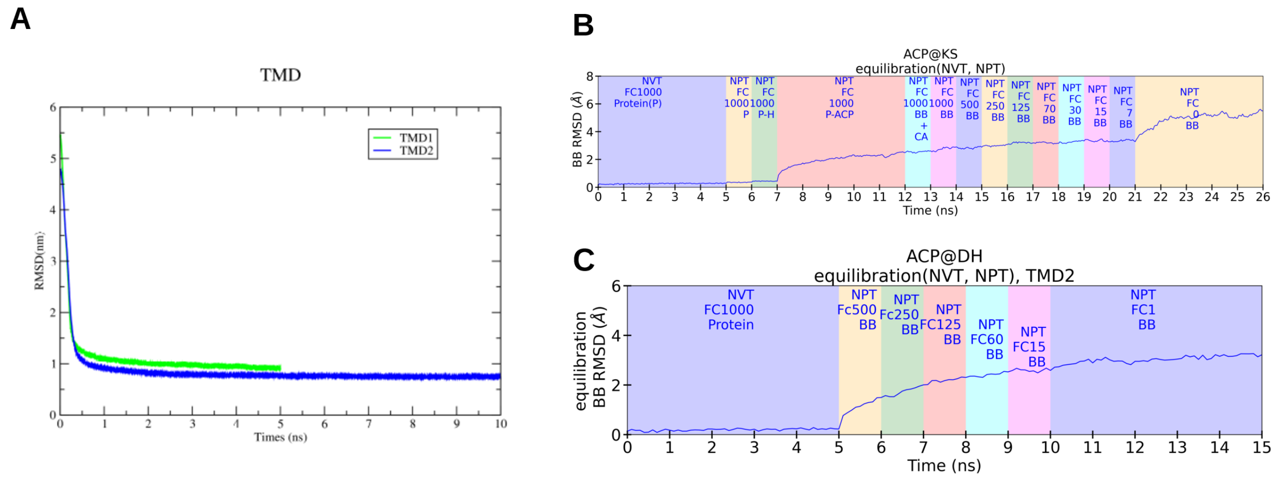


**Figure S6. RMSD for TMD1, TMD2 and slow equilibration used in traj1$_{50ns}$ and traj2$_{100ns}$ (related to section 7.3 and 7.4 ).** (A) RMSD for Cα of ACP as a function of time shows gradual decrease for both TMD1 (green) and TMD2 (blue) (B) RMSD for backbone atoms with respect to energy minimized structure for equilibration followed during the ACP stalled at KS. (C) RMSD for backbone atoms with respect to energy minimized structure for equilibration followed during TMD2 (ACP stalled at DH). Force Constant (kJ $mol^{-1}$ $nm^{-1}$) are mentioned in figure legend.

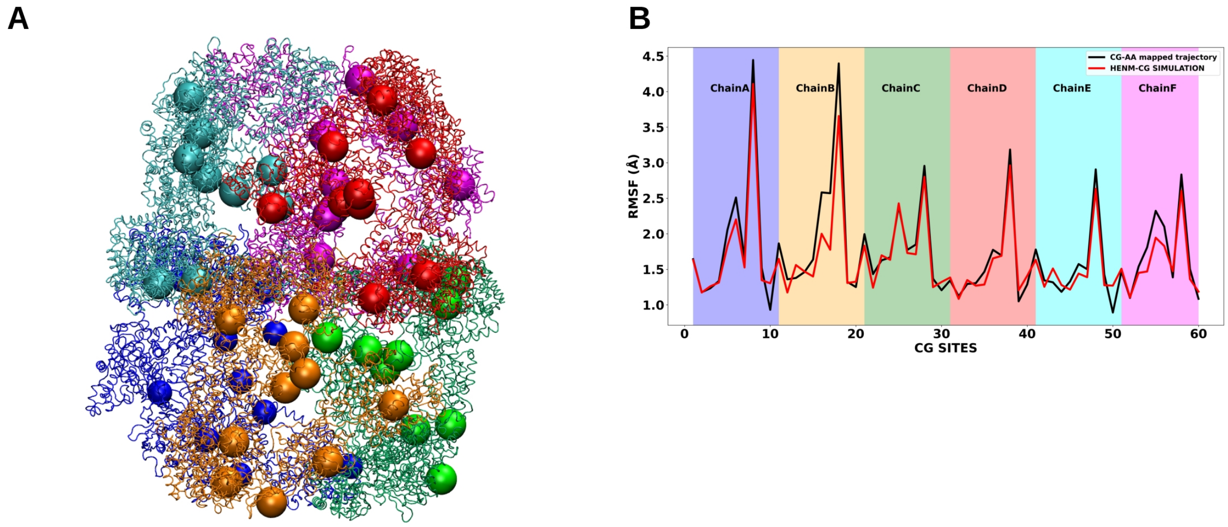


**Figure S7. Coarse grained HENM model (related to Figure 7).** (A) Coarse-grained sites (colored spheres) of each domain in all six chains of Mtb FAS-I complex, (B) RMSF of target CG-AA mapped trajectory and HENM-CG trajectory.

**Table S1. RMSF of seven catalytic domain and linkers for all six chains in traj2$_{100ns}$(related to Figure 3).**

| | **AT** | **ER** | **DH** | **MPT** | **MPT (1451:1498)** | **PL** | **ACP** | **CL** | **KR** |
|---|---|---|---|---|---|---|---|---|---|
| **Chain A** | 1.74 ± 0.4 | 1.37 ± 0.4 | 1.74 ± 0.4 | 2.15 ± 0.7 | 2.82 ± 0.8 | 5.56 ± 1.2 | 2.58 ± 1.1 | 6.19 ± 1.5 | 2.26 ± 1.2 |
| **Chain B** | 2.19 ± 0.6 | 1.57 ± 0.5 | 2.4 ± 0.3 | 2.25 ± 0.8 | 3.08 ± 1.1 | 4.27 ± 1.6 | 3.67 ± 0.9 | 5.91 ± 1.4 | 1.91 ± 0.8 |
| **Chain C** | 2.16 ± 0.6 | 1.87 ± 0.4 | 2.62 ± 0.5 | 3.51 ± 0.7 | 3.58 ± 0.8 | 4.43 ± 1.8 | 3.84 ± 1.3 | 7.14 ± 1.5 | 2.4 ± 0.8 |
| **Chain D** | 2.24 ± 0.4 | 1.54 ± 0.4 | 1.95 ± 0.4 | 2.37 ± 0.9 | 3.51 ± 0.8 | 4.78 ± 1.0 | 3.67 ± 0.9 | 6.56 ± 1.6 | 2.08 ± 1.3 |
| **Chain E** | 2.76 ± 0.6 | 1.68 ± 0.4 | 2.14 ± 0.4 | 2.4 ± 0.9 | 3.53 ± 1.0 | 4.68 ± 1.6 | 3.57 ± 1.2 | 7.47 ± 2.2 | 2.12 ± 1.1 |
| **ChainF** | 2.35 ± 0.4 | 1.63 ± 0.6 | 2.37 ± 0.7 | 3.99 ± 1.2 | 5.82 ± 1.3 | 4.51 ± 1.5 | 2.97 ± 0.7 | 6.14 ± 0.7 | 2.62 ± 1.1 |
| **Average for six chains** | 2.24 | 1.61 | 2.2 | 2.61 | 3.72 | **4.71** | **3.38** | **6.57** | 2.23 |

**Table S2. Force constant used in TMD1 (RMSD of Cα atoms of ACP alone) and TMD2 (RMSD of Cα atoms of ACP and PL) while shifting ACP from KS to DH (related to section 7.3 and 7.4).**

| TMD1 (5ns) | | | TMD2 (10ns) | | |
|---|---|---|---|---|---|
| **Step** | **RMSD (nm)** | **Kappa** | **Step** | **RMSD (nm)** | **Kappa** |
| 0 | 5.5 | 0 | 0 | 4.8 | 0 |
| 10000 | 5 | 10000 | 10000 | 4.5 | 10000 |
| 20000 | 4.5 | 10000 | 20000 | 4.25 | 10000 |
| 30000 | 4 | 10000 | 30000 | 4 | 10000 |
| 40000 | 3.5 | 10000 | 40000 | 3.5 | 10000 |
| 50000 | 3 | 10000 | 50000 | 3 | 10000 |
| 60000 | 2.5 | 10000 | 70000 | 2.5 | 10000 |
| 70000 | 2 | 10000 | 80000 | 2 | 10000 |
| 80000 | 1.5 | 10000 | 90000 | 1.5 | 10000 |
| 90000 | 1 | 10000 | 90000 | 1 | 10000 |
| 100000 | 0.5 | 10000 | 100000 | 0.5 | 10000 |
| | | | 110000 | 0 | 10000 |
| | | | 120000 | 0 | 20000 |
| | | | 130000 | 0 | 30000 |
| | | | 140000 | 0 | 42500 |
| | | | 150000 | 0 | 55000 |
| | | | 160000 | 0 | 67500 |
| | | | 170000 | 0 | 70000 |

* nanometer (nm), *Target molecular dynamics simulation (TMD)

**Video S1: Positional characterization of ACP and PL in Mtb FAS-I complex near KS and DH.**

**Video S2: Mtb FAS-I PL unique conformation in chain D in TMD1.**

**Video S3: Structure of Mtb FAS-I ACP in last 25 ns for all chains in traj$_{100ns}$ and α-3 helix conformation.**

**Video S4: ACP recognition helix residue SER1787 is in catalysis range of DH.**

**Video S5: Structure core interaction with DH and PL adsorb on ACP in chain B.**

### References


1. Colizzi, F., Recanatini, M., and Cavalli, A. (2008). Mechanical Features of Plasmodium falciparum Acyl Carrier Protein in the Delivery of Substrates. J. Chem. Inf. Model. *48*, 2289–2293. 10.1021/ci800297v.
